# Profiling ranked list enrichment scoring in sparse data elucidates algorithmic tradeoffs

**DOI:** 10.1101/2024.06.03.597180

**Authors:** Alexander T. Wenzel, John Jun, Ted Liefeld, Pablo Tamayo, Jill P. Mesirov

## Abstract

Gene Set Enrichment Analysis (GSEA) is a method for quantifying pathway and process activation in groups of samples, and its single sample version (ssGSEA) scores activation using mRNA abundance in a single sample. GSEA and ssGSEA were developed for “bulk” samples rather than individual cell technologies such as microarrays and bulk RNA-sequencing (RNA-seq) data. The growing use of single cell RNA-sequencing (scRNA-seq) raises the possibility of using ssGSEA to quantify pathway and process activation in individual cells. However, scRNA-seq data is much sparser than RNA-seq data. Here we show that ssGSEA as designed for bulk data is subject to some amount of score uncertainty and other technical issues when applied to individual cells from scRNA-seq data. We also show that a ssGSEA can be applied robustly to “pseudobulk” aggregate groups of a few hundred to a few thousand cells provided appropriate normalization is used. Finally, in comparing this approach to other ranked list enrichment methods, we find that the UCell method is most robust to sparsity. We have made the aggregate cell version of ssGSEA available as a Python package and GenePattern module and will also modularize UCell for use on GenePattern as well.

## Introduction

The advent of scRNA-seq has enabled the profiling of gene expression in individual cells, a major advantage over past measurement of “bulk” samples which may mask inherent heterogeneity. scRNA-seq data can be used to quantify the abundance of fine grain cell types within a sample, giving rise to computational methods capable of labeling [1,2] and modeling the interactions between these cell types [3]. Furthermore, with emerging spatial imaging technologies accompanying scRNA-seq, the expression patterns within individual cells can be interpreted in the context of their in situ arrangement and distribution, enabling a better understanding of healthy and diseased tissue function [4].

In the field of computational method development for scRNA-seq data, existing techniques for analyzing bulk data are often revisited to decide if they can be used “as-is”, adapted to accommodate the differences between bulk and scRNA-seq data, or discarded in favor of new approaches [5]. Among the most prominent differences between scRNA-seq, and a common cause of method incompatibility, is the sparsity of scRNA-seq data. The advantage of mapping RNA molecules back to their cell of origin comes with the drawback of substantially less material per observation (cell) than is available for a bulk RNA-seq observation (sampled admixture of cells). Therefore, computational methods applied to scRNA-seq data must accommodate a data distribution that is much sparser and includes fewer detected expressed genes per cell than bulk data [5].

Among the methods that may have applicability to scRNA-seq data is Gene Set Enrichment Analysis (GSEA) [6,7]. GSEA, as applied to bulk gene expression, first ranks all of the genes in the transcriptome by their differential expression between two classes of samples. It then scores pre-existing gene sets by quantifying the degree to which gene set genes are non-uniformly distributed towards the top or bottom of the ranked list. While GSEA needs two groups of samples to compute a relative pathway enrichment score in one group compared to the other, single sample GSEA (ssGSEA) requires only a single sample and builds its ranked list based on the actual expression of genes in that sample [8]. When considering the problem of measuring pathway activity in single cell data, it appears attractive at first glance to apply ssGSEA to single cells, treating each one as a sample. In fact, we have already observed many users attempt this using ssGSEA as implemented in our GenePattern computational environment [9,10]. We therefore sought to understand any issues with the application of ssGSEA to scRNA-seq data and what adaptations might be made to improve its performance with this type of data.

## Results

### Sparse data can cause instability in ssGSEA scores

We first assessed the performance of ssGSEA in six scRNA-seq datasets hosted on the CELLxGENE platform [11–17] (Table 2) under varying conditions. All data were normalized according to the counts per 10,000 reads (CPTT) according to CELLxGENE specifications (Table 1). Datasets where a function other than the natural logarithm appeared to have been used in the CPTT computation were re-normalized. Each dataset was subset to only those genes shared among all six datasets to make enrichment scores comparable across datasets. Ten cells were randomly sampled from each dataset **(Figure 1B)**. Gene sets for enrichment scoring were created by randomly sampling from the shared genes among all six datasets. Fifteen gene sets were created, five with 20 genes, five with 50 genes, and five with 100 genes **(Supplemental Table 2)**. We used two different scoring metrics for evaluating gene set enrichment in a ranked list of genes. Both GSEA and ssGSEA compute a running sum along the ranked list, incrementing when a gene set member gene is encountered and decrementing otherwise. The weighted Kolmogorov-Smirnov (KS) statistic, most used in GSEA, is the maximum point, either positive or negative, along the running sum curve [6,7]. The area under the curve (AUC) statistic is the area under the running sum curve and is most used in ssGSEA [8].

**Table 1:**
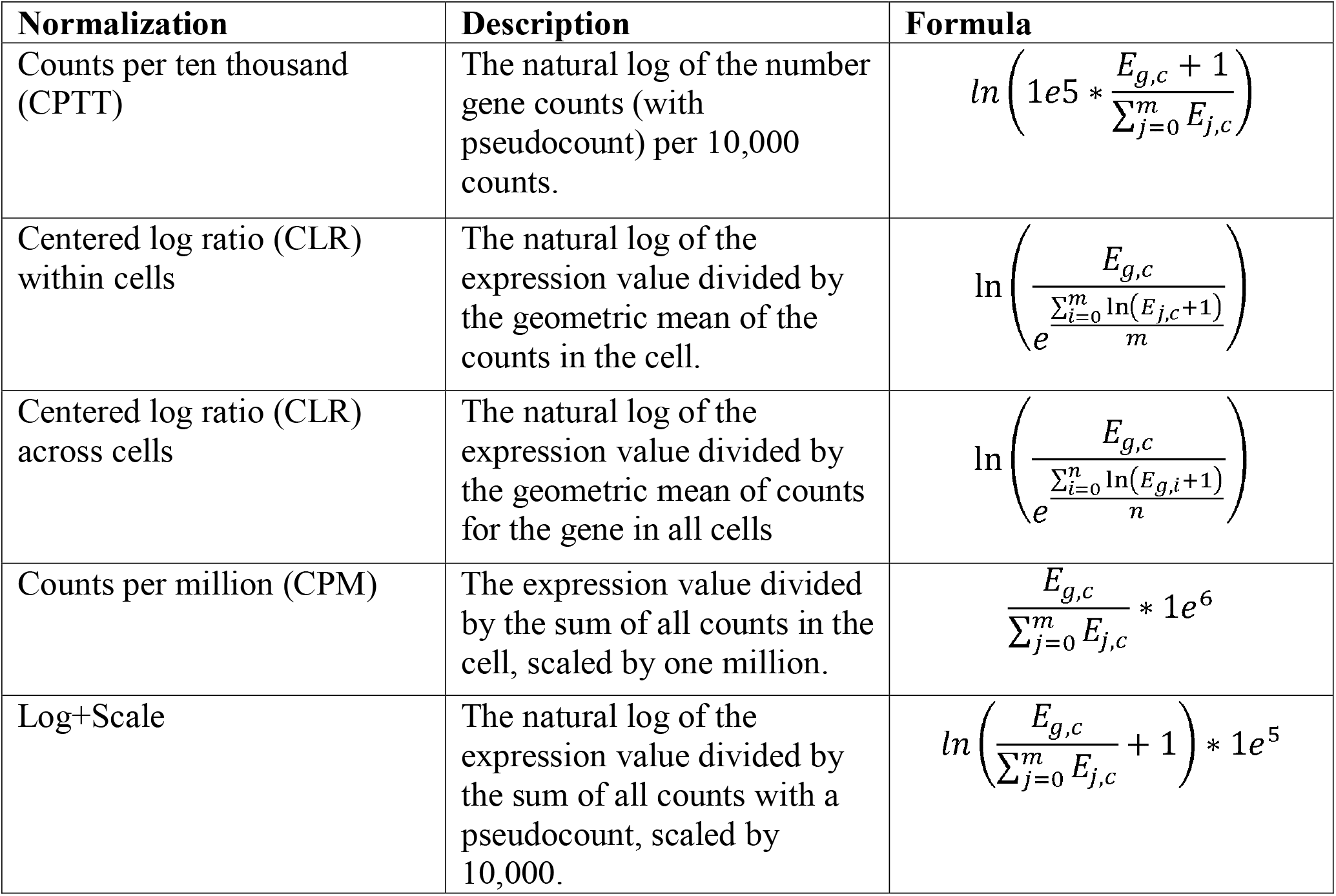
Normalization methods. Normalizations used in benchmarking analysis. For each formula, let *E* be a *m* x *n* scRNA-seq expression matrix of *m* genes and *n* cells. Each formula gives the normalized expression value for the raw counts for gene *g* and cell *c* from *E*.

**Table 2:**
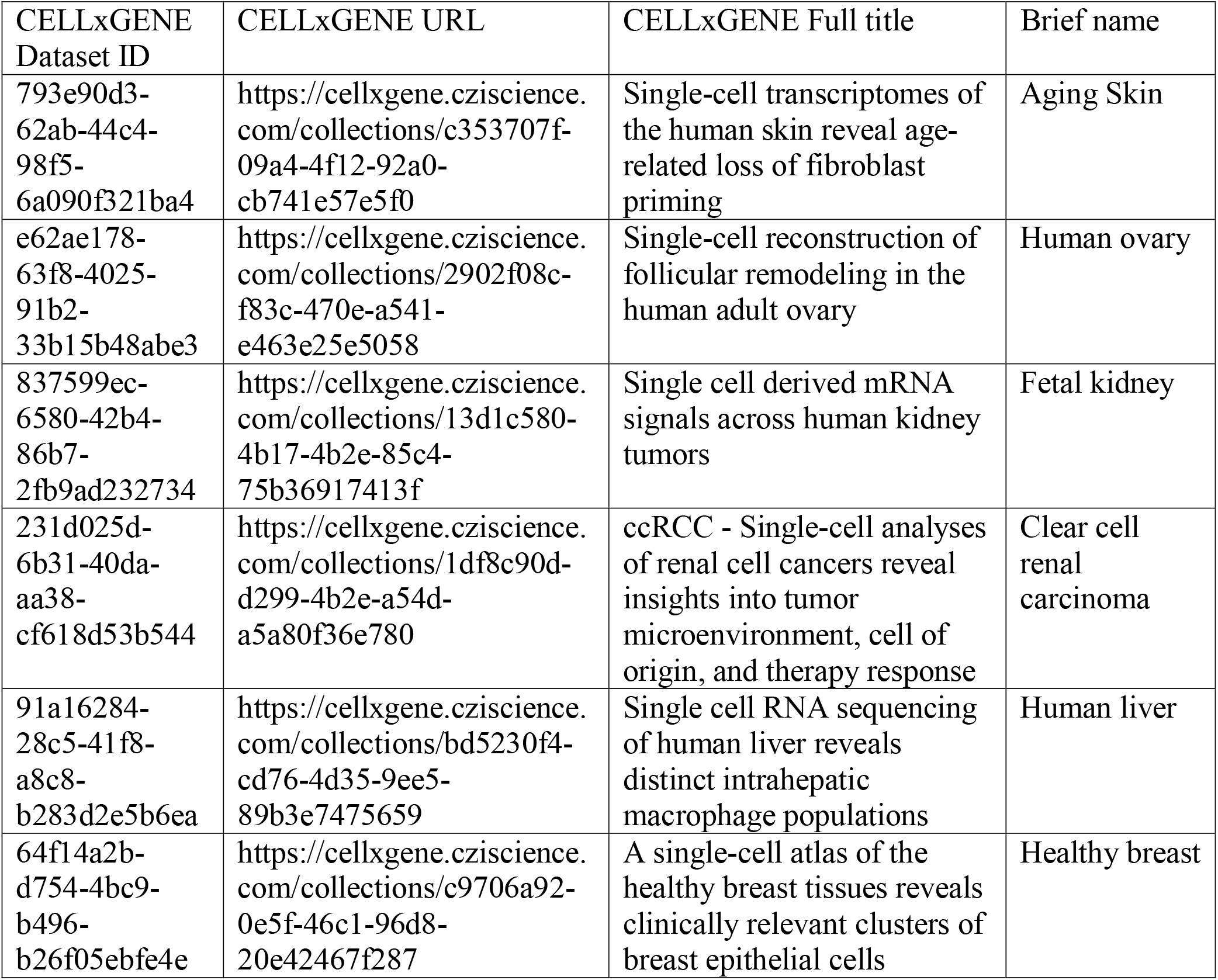
Datasets used in individual and aggregate cell benchmarking. First three columns contain metadata from the CZI CELLxGENE data portal as provided by the authors of the corresponding studies. The fourth column “Brief names” were assigned for the purpose of this work.

**Figure 1:**
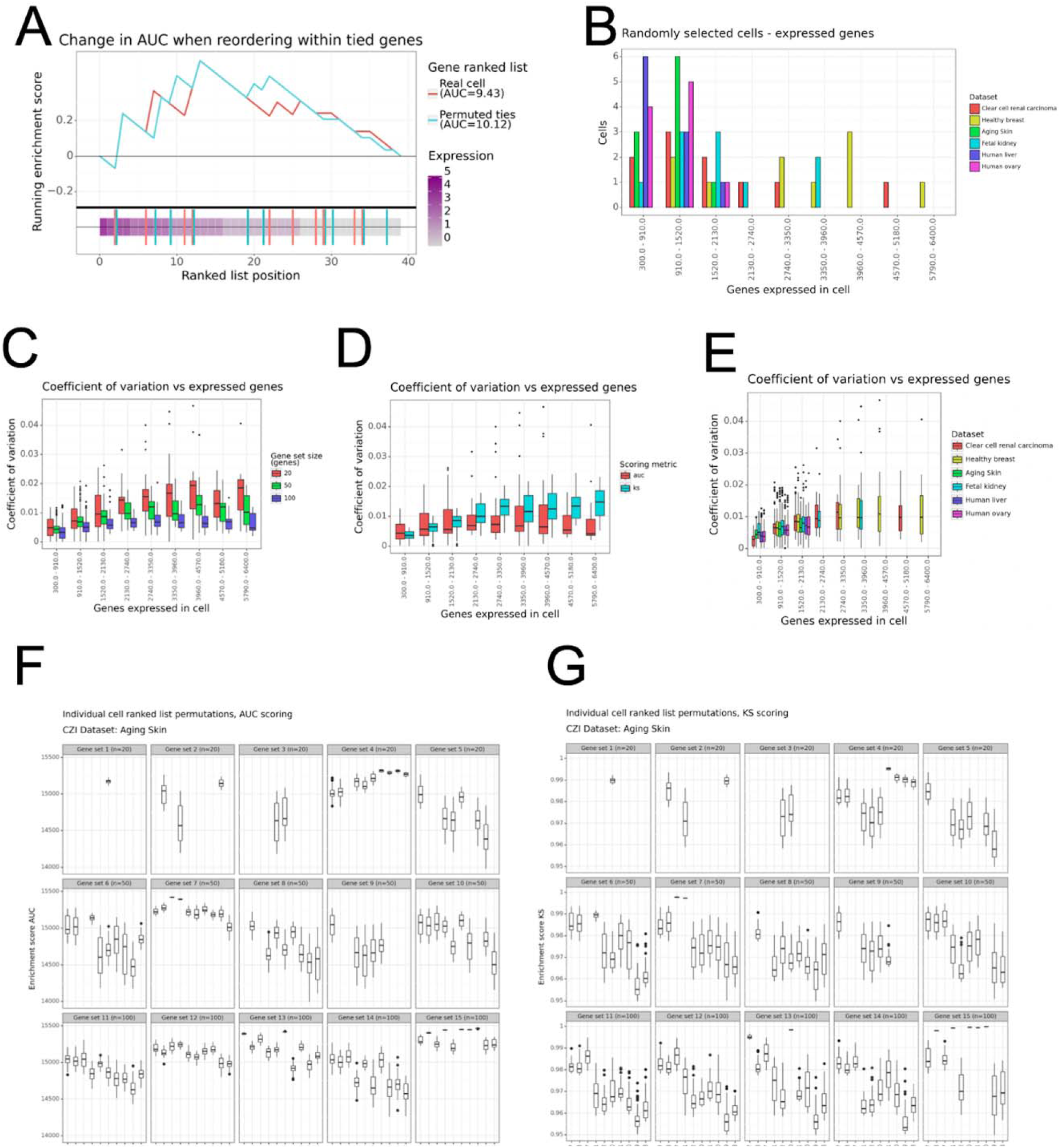
Ranked list permutations of individual cells. **A)** An example of permuting the ranked list of genes in a cell and the resulting ssGSEA scores. Red: GSEA running sum for original list of genes sorted by expression. Blue: GSEA running sum for list of genes where order within genes with equivalent expression has been shuffled. Vertical lines: Gene set gene locations for the original ranked list (Red) and permuted ranked list (Blue). **B)** Distribution of the number of genes expressed in randomly sampled cells. **C-E)** Coefficient of variation for individual cells subjected to ranked list permutations (n = 50) as shown in **(A)**, colored by **(C)** gene set size, **(D)** scoring metric, and **(E)** dataset. **(F-G)** Distribution of enrichment scores in a single dataset “Aging skin”. Each panel is a gene set, each row of panels is a gene set size. Each box represents a single cell and each point represents the enrichment score of a single ranked list permutation of that cell. Scores are separated into **(F)** AUC and **(G)** KS.

To assess the stability of enrichment scores under sparse conditions, we ranked genes based on their expression in each cell and permuted the order of tied genes in the ranked list. Genes are first grouped by their expression value, with all genes having the same expression value placed in the same group. The order of these genes is then shuffled, and the permuted cell is formed by concatenating the shuffled groups. In this benchmarking, we performed 50 such permutations for each of the cells sampled from each dataset. The effect of these permutations is illustrated in the simplified example in **(Figure 1A)**, which demonstrates that it is possible to alter the enrichment score of a sparse sample (cell) with many genes having the same ranking (expression) value without changing the genes’ expression or the gene set used. This is because the reordering of genes within groups of tied genes changes the peaks of the running sum curve, altering the AUC and, in some cases, the KS score. Because GSEA and ssGSEA break ties arbitrarily, both enrichment scores in **(Figure 1A)** are valid, resulting in uncertainty. We measure this uncertainty with the coefficient of variation (CoV), the standard deviation divided by the mean of the enrichment scores for all ranked list permutations, a higher CoV indicates more uncertainty in the enrichment score.

In general, the variance of the CoV itself increased with the number of genes expressed in the cell **(Figure 1C-E)**, as more genes in the gene set being expressed offers more opportunities to perturb the enrichment score by moving gene set genes. Smaller gene sets were more likely to generate higher CoV values **(Figure 1C)**, owing to the greater relative importance of moving one gene within a tied group of genes. AUC scores were more likely to have high AUC values than KS scores in cells with fewer genes expressed **(Figure 1D)**, though we note that there are relatively few cells in our data with more than 3,000 genes expressed **(Figure 1B)**. The KS score depends on the maximum point in the running sum statistic and having fewer gene set genes expressed offers fewer opportunities for perturbing the score. In the process of generating benchmarking, technical issues due to extreme sparsity emerged. If no members of a gene set are expressed at a level above 0, this will result in a division by zero during the GSEA computation for either KS or AUC. For smaller gene sets, some enrichment scores could not be computed **(Figure 1F-G, Supplemental Figure 1-10)**.

For sets where a score could be computed, scores tended to be relatively high **(Figure 1F-G)**, especially when using the KS scoring metric **(Figure 1G)**. In a sparse sample, most of the signal for enrichment comes from the small fraction of strongly expressed genes in a cell. The reduced detection for moderately expressed genes results in more extreme enrichment scores driven by a small number of highly expressed genes. A major advantage of GSEA is the ability to detect collectively upregulated pathways even when no single gene is strongly differentially expressed [6], and this advantage is reduced when computing enrichment on an extremely sparse single cell.

### Adapting ssGSEA to high-sparsity conditions

As the sparsity of individual cells both causes uncertainty and prevents the computation of enrichment scores in some single cells with some gene sets, we concluded that aggregating multiple cells is preferable for using ssGSEA on scRNA-seq data. Here we describe our benchmarking of aggregation via averaging the expression of cells within each pre-annotated cell type in each CELLxGENE dataset. We benchmarked the performance of ssGSEA using four normalization methods **(Table 1)**, as well as the two scoring metrics and 15 gene sets as described above. Each dataset was evaluated using raw counts and normalized with four normalization methods and the gene expression for cells within each annotated cell type were averaged. These aggregate average cells were subjected to the same ranked list permutation as described above.

Representative benchmarking results for one dataset are shown in **Figure 2A** and **Figure 2B** (for remaining datasets, see **Supplemental Figures 2.11-20**). For AUC scoring, we observed relatively little difference between non-normalized counts and normalized counts, indicating that sufficiently large groups of tied genes still exist to create uncertainty in the AUC score after normalization and aggregation **(Figure 2A, Supplemental Figures 2.11-20)**. For KS scoring, most normalized scores had CoVs sufficiently small enough as to be considered effectively stable or constant **(Figure 2B)**. Notably for some cell types, normalized data still resulted in relatively high CoV, usually for smaller gene sets **(Figure 2C)**. For most cell types, the KS scores for unnormalized cells had uniformly higher CoV values than their normalized counterparts except for one cell type that contained 20,000 cells **(Figure 2D)**.The distribution of counts for 20,000 averaged cells may begin to approach the distribution of a bulk RNA-seq sample depending on the transcriptional heterogeneity of the included cells, but a cluster or cell type that large in most contemporary datasets originating from a single experiment is uncommon. While both scoring metrics may yield high CoVs for some cell types and gene sets, all of the most stable scores were derived using the KS metric **(Figure 2E)**. In fact, the combination of KS scoring with the centered-log-ratio (CLR) across cells **(Table 1)** resulted in the most stable scoring, with nearly all combinations of cell type and gene set yielding a CoV less than 10e-5 (**Figure 2F)**. Given these results, we selected KS as the scoring metric and the CLR applied across cells as the normalization method for scGSEA, as that combination yielded the most stable results.

**Figure 2:**
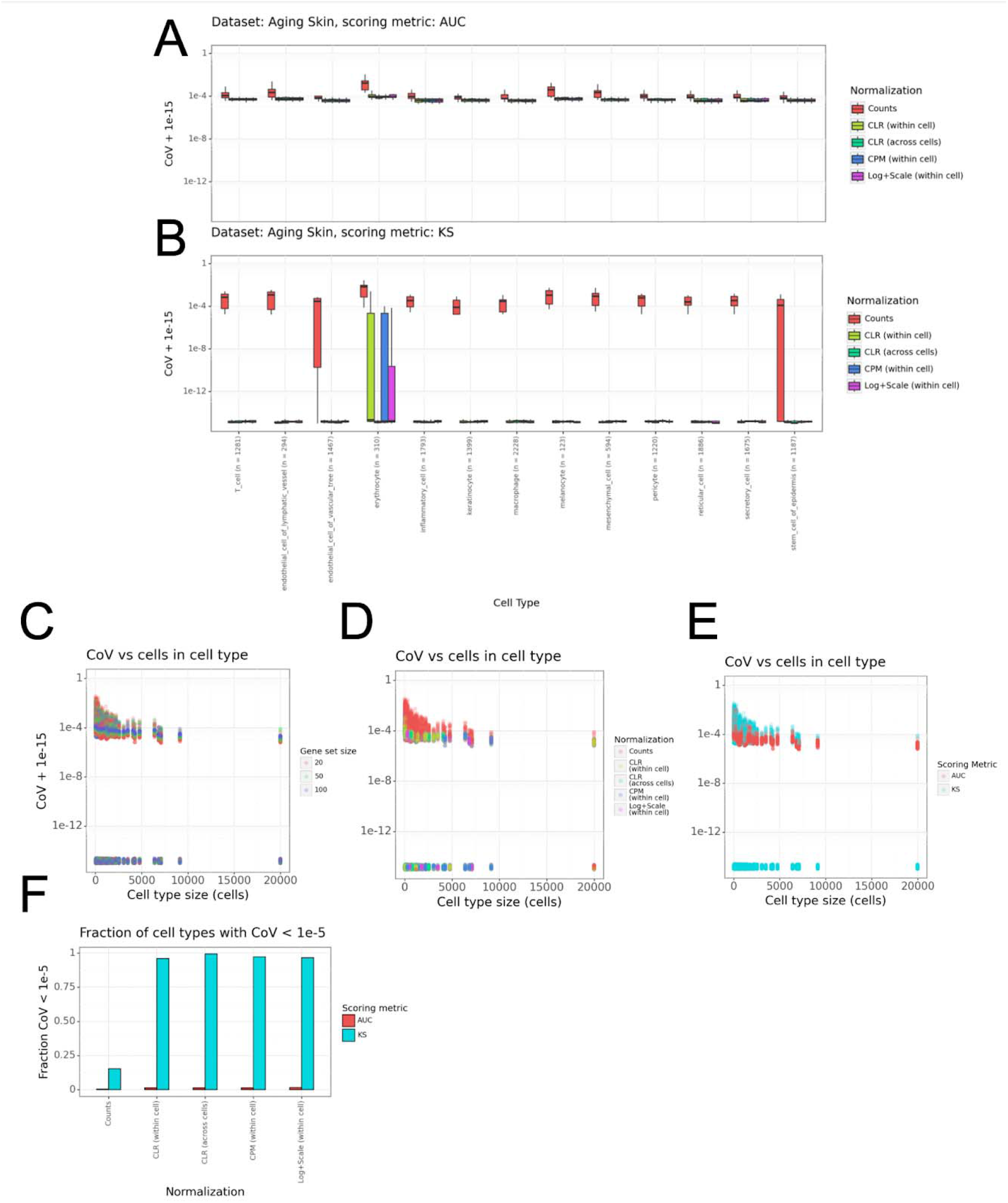
Ranked list permutation of aggregate cells. **A-B)** Distribution of CoV values for each cell type in the “Aging skin” dataset. Each point represents the CoV of enrichment scoring for one gene set on permutations of a ranked list following normalization and averaging of cells in the cell type. Results are separated by **(A)** AUC and **(B)** KS scores. **C-D)** CoV of enrichment scores on ranked list permutations versus number of cells in the given cell type. Each point is the CoV of enrichment scores for a single cell type, gene set, normalization, and scoring metric. Points are colored by **(C)** gene set size, **(D)** normalization method, and **(E)** scoring metric. **F)** Fraction of CoV values below 1e-5 for each choice of normalization method and scoring metric.

### ssGSEA on aggregate cells reduces uncertainty to separate cell types using expression signatures

As an application of scGSEA, we turned to the problem of separating cell types based on pathway enrichment scores. We obtained two datasets representing two immune cell populations in the Tabula Sapiens atlas (The Tabula Sapiens Consortium, 2022) as distributed in CELLxGENE [11]: spleen-resident B cells and bone marrow-resident plasma cells **(Figure 3A)**. We selected two experimentally derived gene sets [18] from MSigDB [19] representing spleen-resident B cells and bone marrow-resident plasma cells respectively. We first used the currently available ssGSEA method in GenePattern [9] to score the two gene sets in each of the individual B cells and plasma cells **(Figure 3B)**. In general, the B cell gene set scored higher in the B cells and the plasma cell gene set scored slightly higher in the plasma cells, but with a wide range of scores in both cases **(Figure 3B)**. To understand the role of instability, we selected ten cells at random from each group and subjected them to the ranked list permutation approach described above. **Figure 3C** shows the range of scores obtained in each cell. Notably, the range of valid scores overlapped in many cases for pairs of cells from the two distinct populations, meaning that sparsity and the resulting uncertainty prevents the classification of individual cells as B cells or plasma cells based on cell type signature enrichment using ssGSEA.

**Figure 3:**
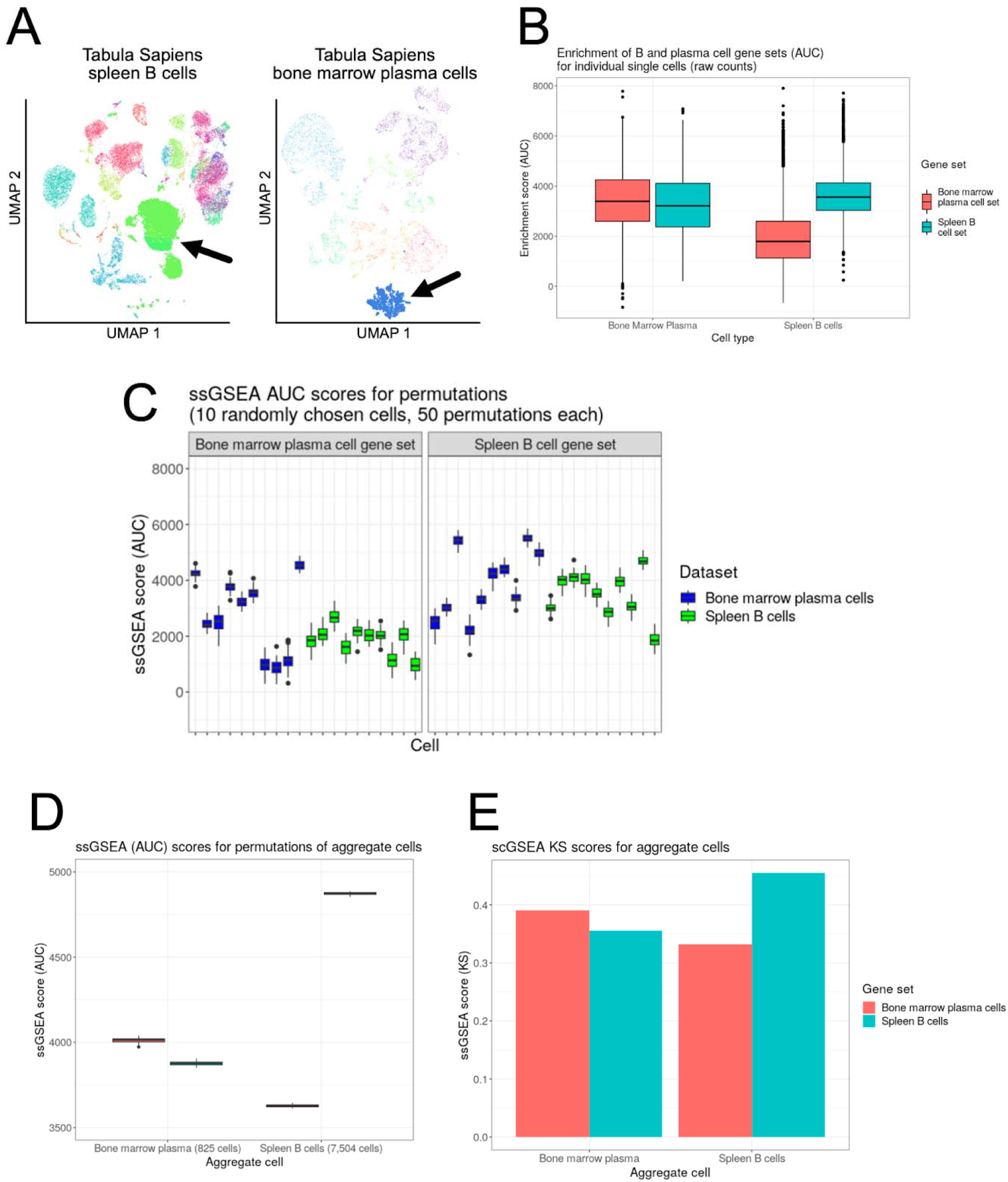
scGSEA separates B cells and plasma cells with more certainty. **A)** CELLxGENE-hosted datasets for Tabula Sapiens spleen cells (left) and bone marrow cells (right). UMAP plots are as rendered by the CELLxGENE data explorer. Arrows point to spleen B cells (lime green, left) and bone marrow plasma cells (blue, right). **B)** Enrichment scores for the spleen B cell gene set (teal) and bone marrow plasma cell gene set (teal) scores in the Tabula Sapiens bone marrow plasma cells (left) and spleen B cells (right). **C)** Scores for 10 randomly selected cells from the two Tabula Sapiens datasets following ranked list permutations (n = 50) applied to each cell. **D-E)** Scores for ranked list permutations (n = 50) following CLR normalization and averaging of cells within the spleen B cell group (teal) and bone marrow plasma cell group (salmon). Scores are split into **(D)** AUC and **(E)** KS. **(E)** is rendered as a bar plot because all scores were the same across permutations.

We then used the scGSEA approach as described above, normalizing with the CLR across cells and creating an aggregate cell for each cell type. We then applied the same ranked list permutation approach to the two aggregate cells. Though we selected the KS score as the scoring metric based on scGSEA, we also evaluated the AUC as the scoring metric here. Though the uncertainty of AUC scores was greatly reduced **Figure 3D**, the uncertainty was effectively eliminated with the KS score, as we had observed in the benchmarking (**Figure 3E**). Using this combination of normalization and KS scoring, we were able to separate the bone marrow plasma cell group from the spleen B cell group.

### UCell shows most stability among single cell ranked list enrichment methods

We next assessed stability of enrichment scores in other methods using ranked lists of genes for enrichment scoring in GSEA. We evaluated GSVA [20], AUCell [21], and UCell [22]. We used GSVA’s “ssgsea” option, which is similar to our existing ssGSEA method using rank normalization and the AUC as the enrichment score. AUCell computes an enrichment score derived from the area under the empirical cumulative distribution of gene set genes in the ranked list. UCell tests for enrichment with a variation of the Mann-Whitney U test, accounting for the size of the gene set being tested and a user-provided “maxRank” parameter, a cutoff value for the worst-ranked gene that the method will consider.

We applied each method to the same ranked list permutations of single cells as described above. For all methods, we used default parameters without modification where possible. Because AUCell internally calls a sorting function on the ranked list, we re-built the AUCell Python package with this sorting removed so that we could control the order of tied genes in ranked lists externally. This modified AUCell package, as well as the other methods we tested, are installed in a publicly available Docker image (see **Code availability**).

**Figure 4A** shows the distribution of coefficients of variation across the same permuted cells and randomly sampled gene sets as used above. GSVA, using a similar approach to ssGSEA’s AUC, had the widest variation in scores, in line with the results of ssGSEA itself. While AUCell had a small amount of instability in some cases, UCell produced the exact same score in all cases. This is because UCell by default assigns the same rank value to each tied gene and, because the score is derived from summing the ranks of gene set genes, the order within those tied groups should not alter the score. To ensure that the lack of variability was not due to UCell’s maxRank cutoff (1500 by default), we ran this analysis with maxRank = 3000 and maxRank = 5000. This additional profiling did not produce a coefficient of variation above 1e-32, a negligible amount most likely attributable to machine precision. We observed that UCell’s maxRank cutoff default value is relatively stringent, excluding genes with expression values as high as 4 UMI counts (**Figure 4B**). In our benchmarking, a maxRank of 5000 was sufficient to only exclude genes with 1 count or fewer **(Figure 4B)**. Finally, we applied UCell to the same B cell and plasma cell separation task. The distribution of scores on individual cells was similar to those obtained by ssGSEA, but without the accompanying uncertainty in scoring **(Figure 4C)**.

**Figure 4:**
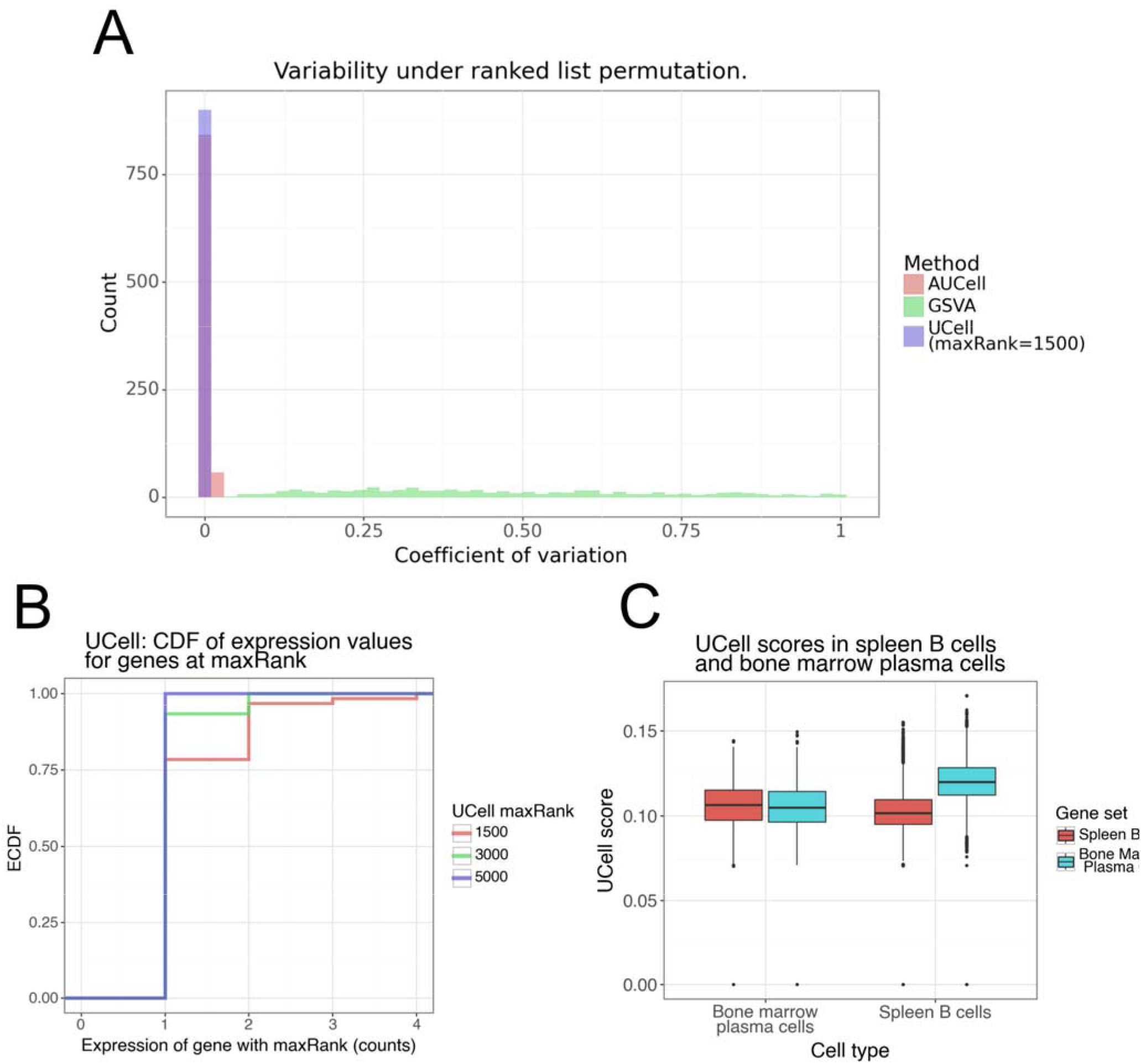
Profiling single cell ranked list enrichment methods. **A)** Coefficient of variation distribution across cell ranked list permutations for the three methods tested. A small number of extreme outliers above 1.0 were excluded. **B)** Cumulative distribution of the expression values for genes with rank values of 1500, 3000, and 5000 in UCell. **C)** Enrichment scores using UCell for bone marrow resident plasma cells and spleen resident B cells using the same data and gene sets as in **Figure 3B**.

## Discussion

To summarize, we highlight a potential pitfall of ranked list enrichment methods when used with sparse single cell data, show that applying ssGSEA specifically to aggregate cells yields stable scores, and identify UCell as the most robust method with respect to enrichment score stability. While the choice to aggregate cells is necessary to avoid uncertainty and cases where there is in fact no valid enrichment score for a given cell and gene set, this aggregation does not significantly reduce scRNA-seq’s ability to yield more granular observations than are obtainable at the bulk level. Because the analysis of bulk data requires the often-oversimplifying assumption that a bulk sample is a homogeneous collection of cells, generating enrichment scores in different subtypes of cells is still a major advantage, even if all of the cells within each cell type must be averaged. However, we do note that our aggregate cell enrichment method does rely on a relative normalization applied to each gene across cells. The authors of UCell highlight this issue in comparing their method to the module score approach implemented in the Seurat toolkit [23], noting that the enrichment score changes if cells are added or removed from the dataset. We find that the UCell approach both solves this issue and yields an enrichment score that is robust to the phenomenon of large groups of ties when creating ranked lists from sparse data. We have made our aggregate cell ssGSEA approach available in our GenePattern cloud platform for accessible, reproducible bioinformatics [9], and we are currently working to deploy UCell as a GenePattern module as well.

## Code availability

The environment for running benchmarking analysis is available on Docker Hub at the tag atwenzel/scgsea-method-comp:v0.2.0. sc-ssGSEA is available as the sc_ssGSEA Python package on PyPI and in a module of the same name on the public GenePattern Cloud server (cloud.genepattern.org).

## References

1. Nofech-Mozes, I., Soave, D., Awadalla, P. & Abelson, S. Pan-cancer classification of single cells in the tumour microenvironment. Nat Commun 14, 1615 (2023).

2. Domínguez Conde, C. et al. Cross-tissue immune cell analysis reveals tissue-specific features in humans. Science 376, eabl5197 (2022).

3. Armingol, E., Officer, A., Harismendy, O. & Lewis, N. E. Deciphering cell–cell interactions and communication from gene expression. Nat Rev Genet 22, 71–88 (2021).

4. Longo, S. K., Guo, M. G., Ji, A. L. & Khavari, P. A. Integrating single-cell and spatial transcriptomics to elucidate intercellular tissue dynamics. Nat Rev Genet 22, 627–644 (2021).

5. Chen, G., Ning, B. & Shi, T. Single-Cell RNA-Seq Technologies and Related Computational Data Analysis. Frontiers in Genetics 10, (2019).

6. Mootha, V. K. et al. PGC-1α-responsive genes involved in oxidative phosphorylation are coordinately downregulated in human diabetes. Nature Genetics 34, 267–273 (2003).

7. Subramanian, A. et al. Gene set enrichment analysis: a knowledge-based approach for interpreting genome-wide expression profiles. Proceedings of the National Academy of Sciences 102, 15545–15550 (2005).

8. Barbie, D. A. et al. Systematic RNA interference reveals that oncogenic KRAS-driven cancers require TBK1. Nature 462, 108–112 (2009).

9. Reich, M. et al. GenePattern 2.0. Nature Genetics 38, 500 (2006).

10. Reich, M. et al. The GenePattern Notebook Environment. Cell Systems 5, 149-151.e1 (2017).

11. Program, C. S.-C. B. et al. CZ CELL×GENE Discover: A single-cell data platform for scalable exploration, analysis and modeling of aggregated data. 2023.10.30.563174 Preprint at 10.1101/2023.10.30.563174 (2023).

12. Bhat-Nakshatri, P. et al. A single-cell atlas of the healthy breast tissues reveals clinically relevant clusters of breast epithelial cells. Cell Reports Medicine 2, 100219 (2021).

13. MacParland, S. A. et al. Single cell RNA sequencing of human liver reveals distinct intrahepatic macrophage populations. Nat Commun 9, 4383 (2018).

14. Zhang, Y. et al. Single-cell analyses of renal cell cancers reveal insights into tumor microenvironment, cell of origin, and therapy response. Proceedings of the National Academy of Sciences 118, e2103240118 (2021).

15. Young, M. D. et al. Single cell derived mRNA signals across human kidney tumors. Nat Commun 12, 3896 (2021).

16. Fan, X. et al. Single-cell reconstruction of follicular remodeling in the human adult ovary. Nat Commun 10, 3164 (2019).

17. Solé-Boldo, L. et al. Single-cell transcriptomes of the human skin reveal age-related loss of fibroblast priming. Commun Biol 3, 1–12 (2020).

18. Benson, M. J. et al. Heterogeneous nuclear ribonucleoprotein L-like (hnRNPLL) and elongation factor, RNA polymerase II, 2 (ELL2) are regulators of mRNA processing in plasma cells. Proc Natl Acad Sci U S A 109, 16252–16257 (2012).

19. Liberzon, A. et al. Molecular signatures database (MSigDB) 3.0. Bioinformatics 27, 1739–1740 (2011).

20. Hänzelmann, S., Castelo, R. & Guinney, J. GSVA: gene set variation analysis for microarray and RNA-Seq data. BMC Bioinformatics 14, 7 (2013).

21. Aibar, S. et al. SCENIC: single-cell regulatory network inference and clustering. Nat Methods 14, 1083–1086 (2017).

22. Andreatta, M. & Carmona, S. J. UCell: Robust and scalable single-cell gene signature scoring. Computational and Structural Biotechnology Journal 19, 3796–3798 (2021).

23. Satija, R., Farrell, J. A., Gennert, D., Schier, A. F. & Regev, A. Spatial reconstruction of single-cell gene expression data. Nature Biotechnology 33, 495–502 (2015).

