## Supplemental Information for "Profiling ranked list enrichment scoring in sparse data elucidates algorithmic tradeoffs"

| **Normalization** | **Description** | **Formula** |
| --- | --- | --- |
| Counts per ten thousand (CPTT) | The natural log of the number gene counts (with pseudocount) per 10,000 counts. | $\ln\left( 1e5*\frac{E_{g,c}+1}{\sum_{j=0}^{m} E_{j,c}} \right)$ |
| Centered log ratio (CLR) within cells | The natural log of the expression value divided by the geometric mean of the counts in the cell. | $\ln\left( \frac{E_{g,c}}{e^{\frac{\sum_{i=0}^{m} \ln\left( E_{j,c}+1 \right)}{m}}} \right)$ |
| Centered log ratio (CLR) across cells | The natural log of the expression value divided by the geometric mean of counts for the gene in all cells | $\ln\left( \frac{E_{g,c}}{e^{\frac{\sum_{i=0}^{n} \ln\left( E_{g,i}+1 \right)}{n}}} \right)$ |
| Counts per million (CPM) | The expression value divided by the sum of all counts in the cell, scaled by one million. | $\frac{E_{g,c}}{\sum_{j=0}^{m} E_{j,c}}*1e^{6}$ |
| Log+Scale | The natural log of the expression value divided by the sum of all counts with a pseudocount, scaled by 10,000. | $\ln\left( \frac{E_{g,c}}{\sum_{j=0}^{m} E_{j,c}}+1 \right)*1e^{5}$ |

**Table 1**: Normalization methods

Normalizations used in benchmarking analysis. For each formula, let *E* be a *m* x *n* scRNA-seq expression matrix of *m* genes and *n* cells. Each formula gives the normalized expression value for the raw counts for gene *g* and cell *c* from *E*.

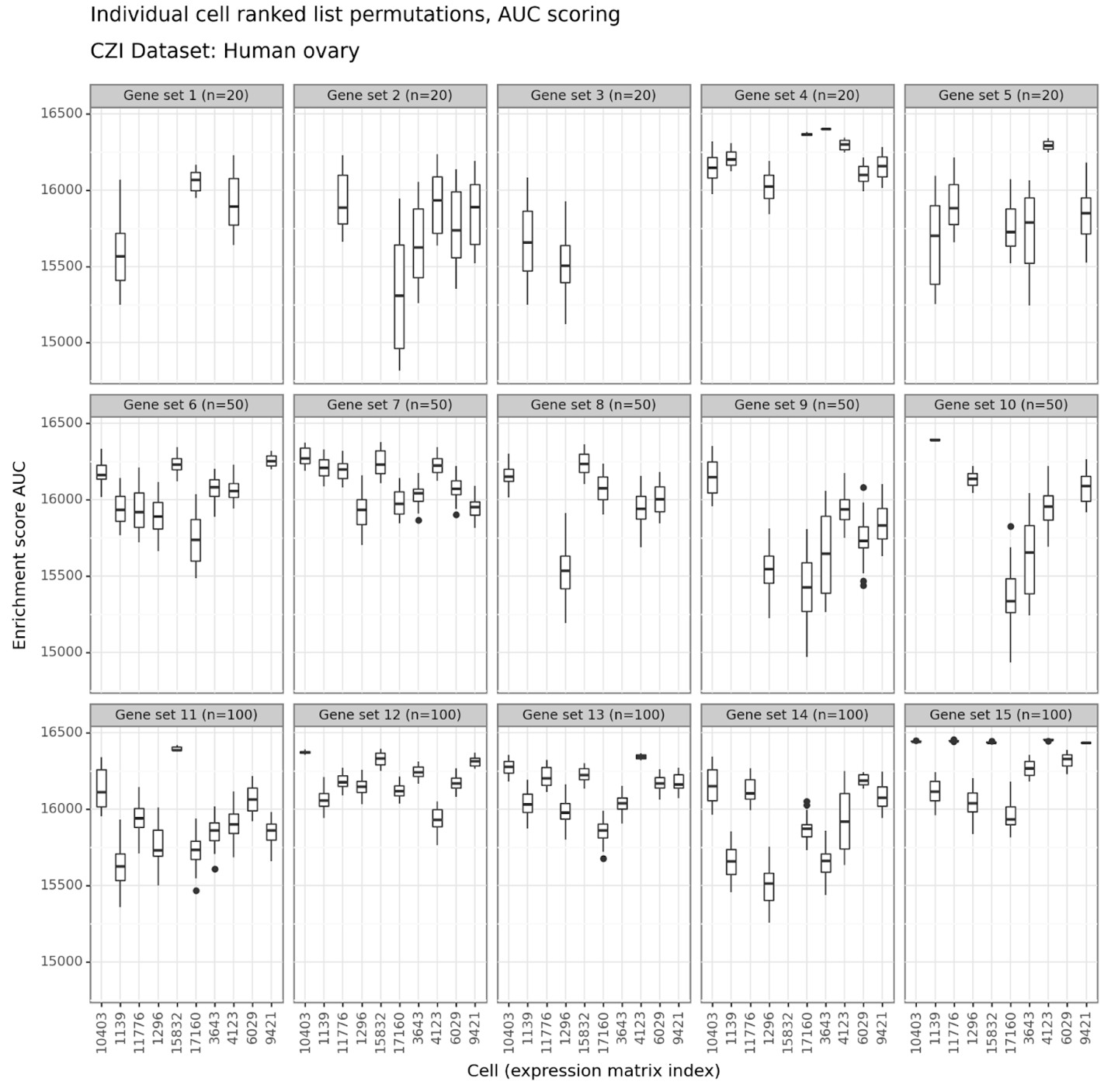

**Figure S.1**: Individual cell permutations – Human ovary – AUC scores

Figure features as in **Figure 1F**

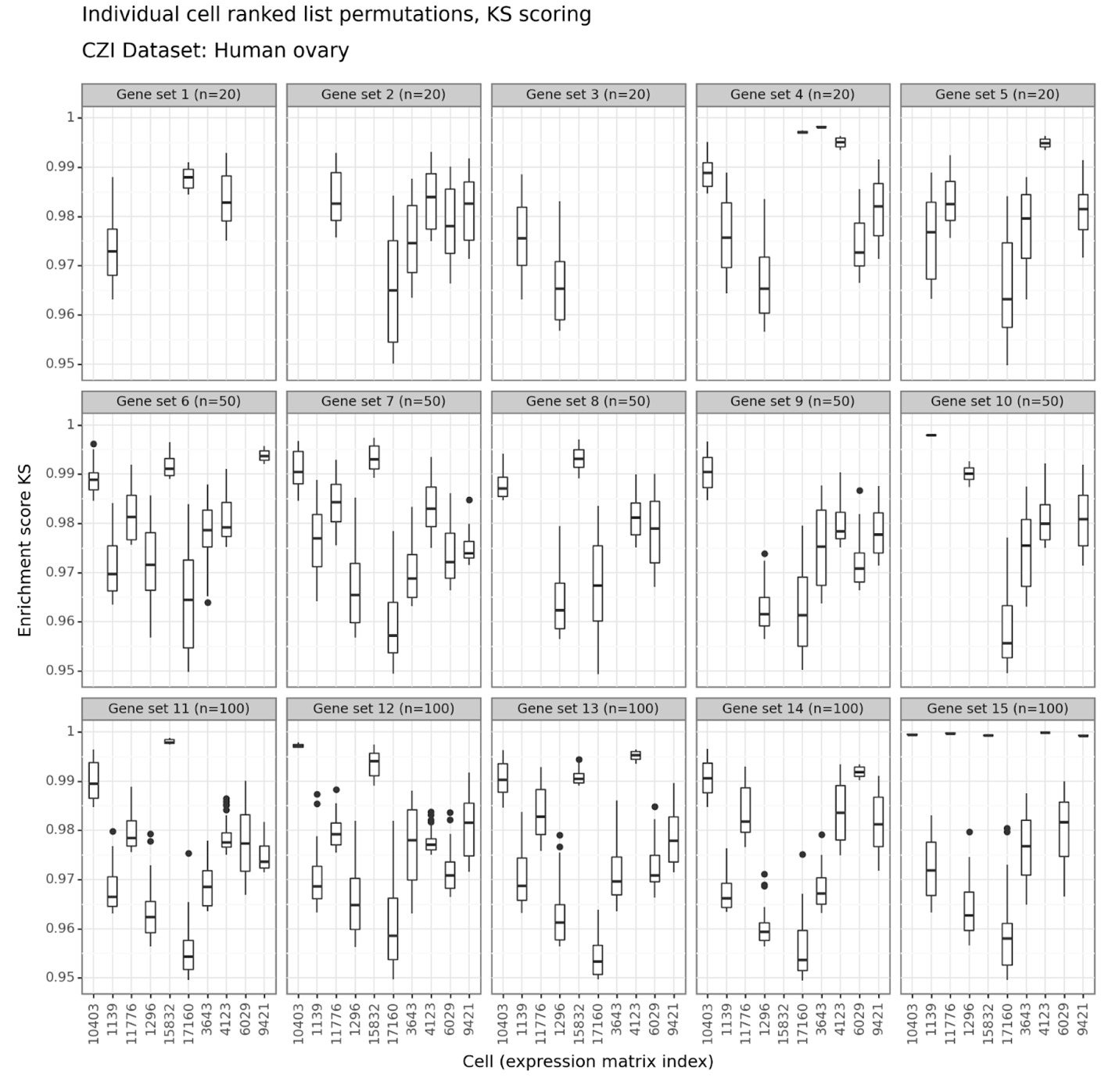

**Figure S2**: Individual cell permutations – Human ovary – KS scores

Figure features as in **Figure 1G**

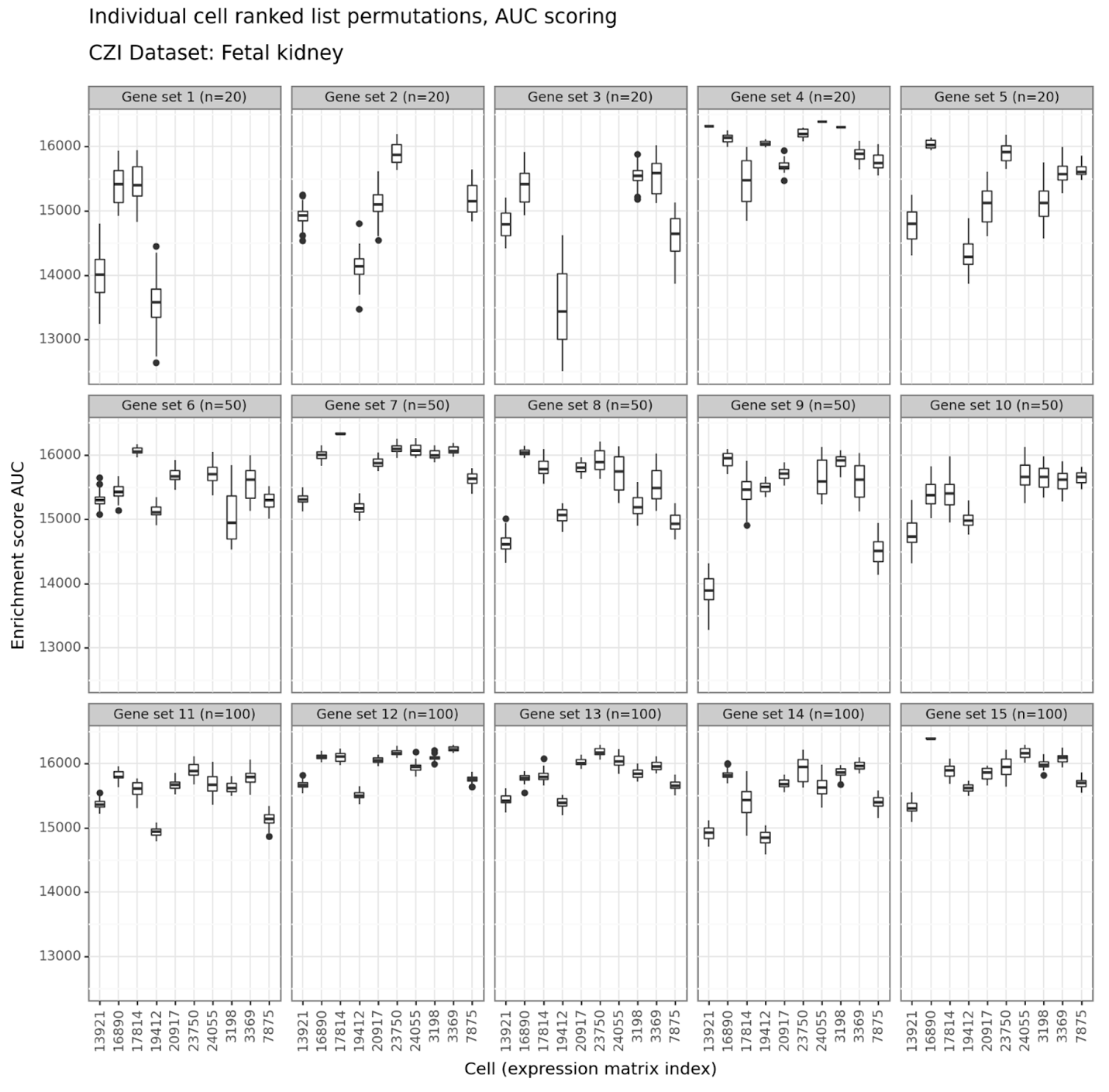

**Figure S3**: Individual cell permutations – Fetal kidney – AUC scores

Figure features as in **Figure 1F**

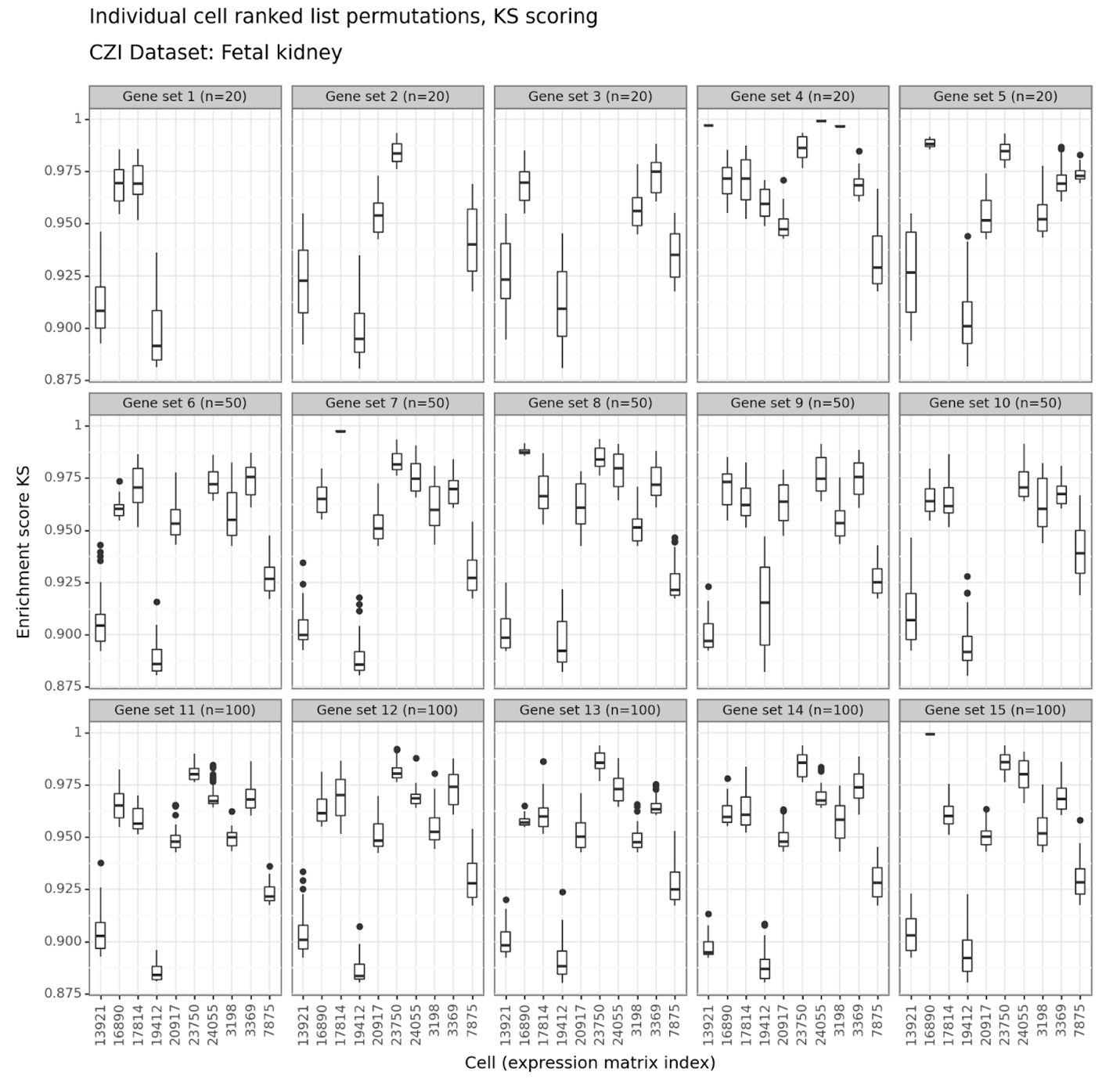

**Figure S4**: Individual cell permutations – Fetal kidney – KS scores

Figure features as in **Figure 1G**

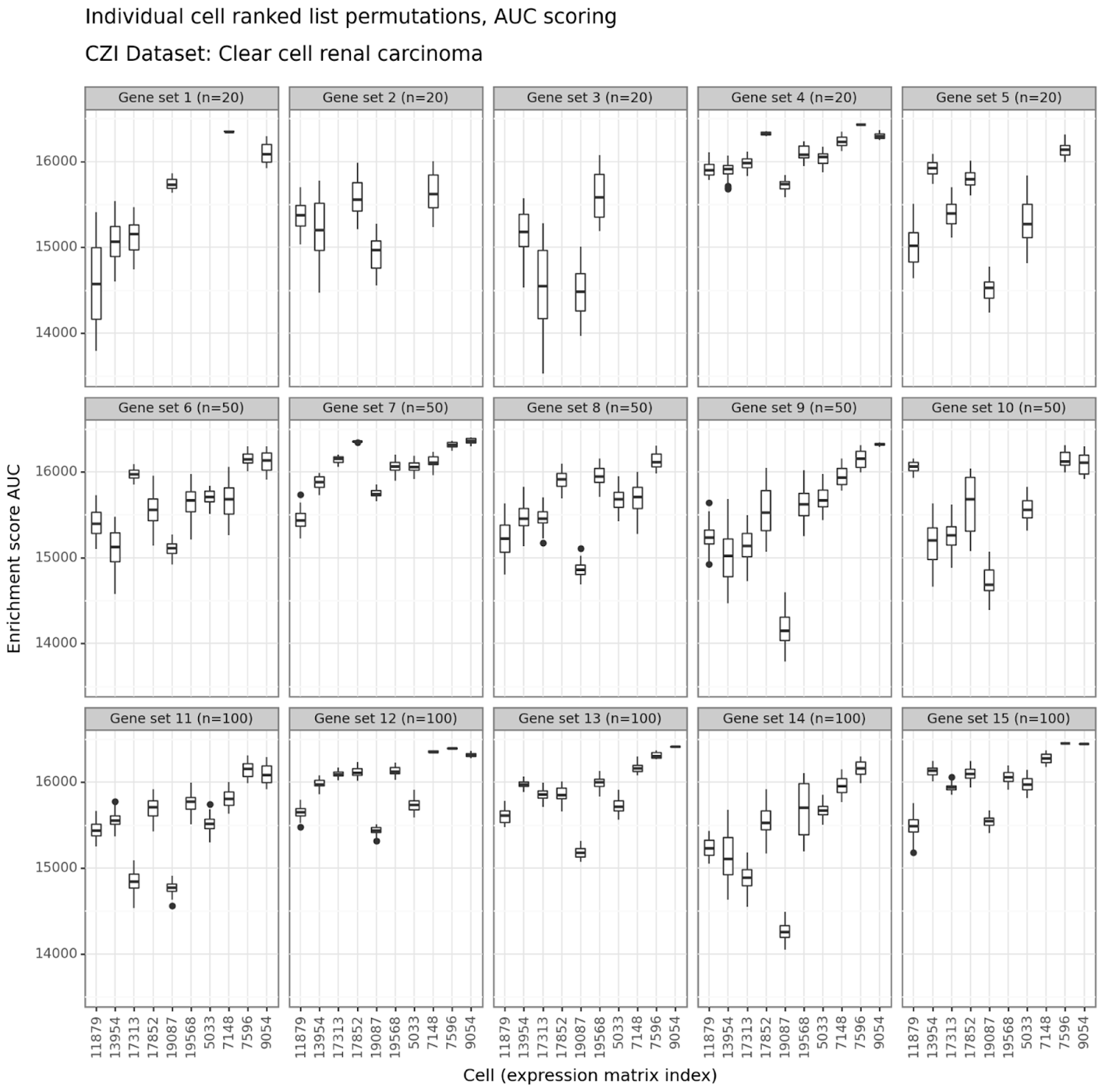

**Figure S5**: Individual cell permutations – Clear cell renal carcinoma – AUC scores

Figure features as in **Figure 1F**

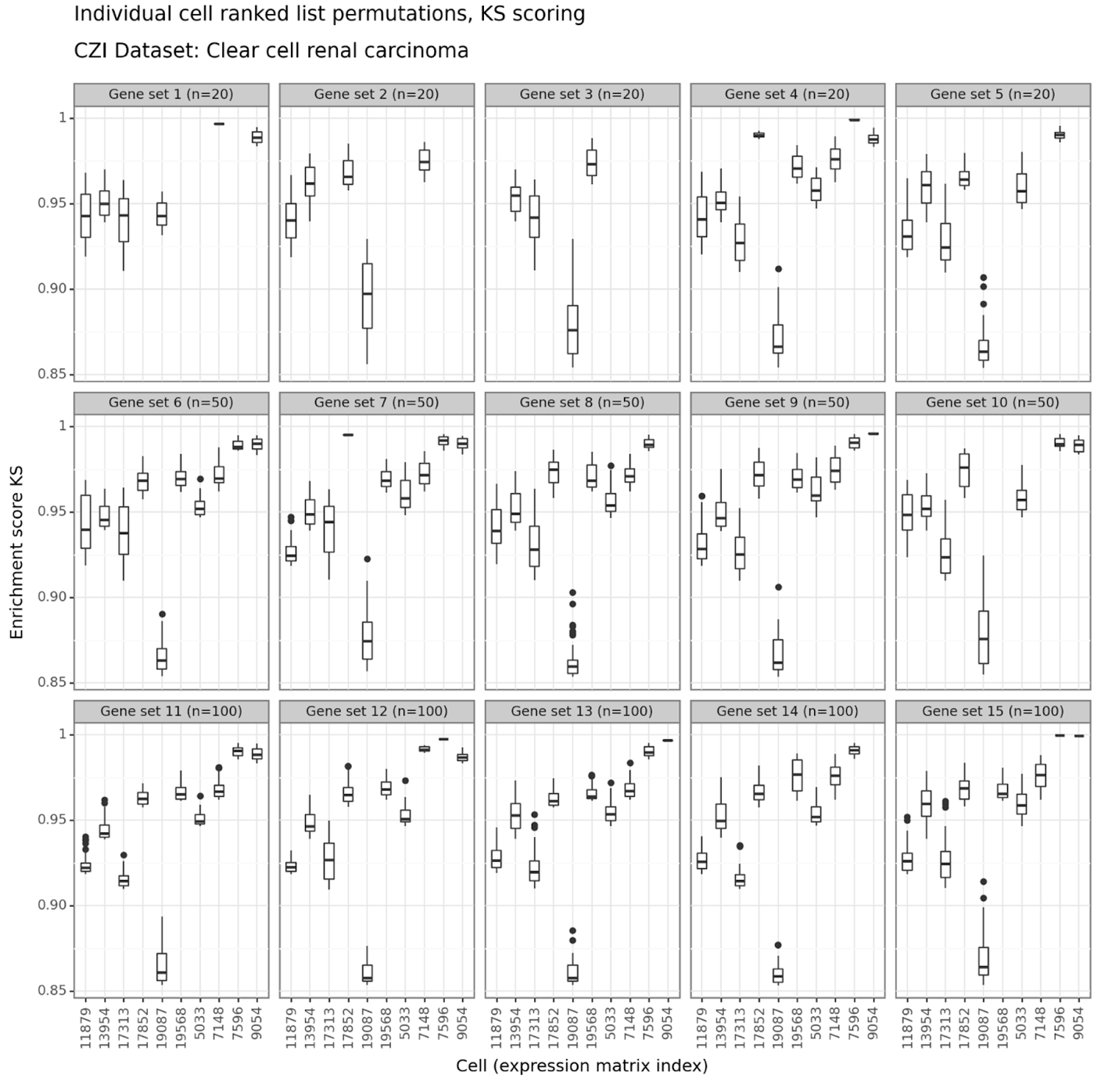

**Figure S6**: Individual cell permutations – Clear cell renal carcinoma – KS scores

Figure features as in **Figure 1G**

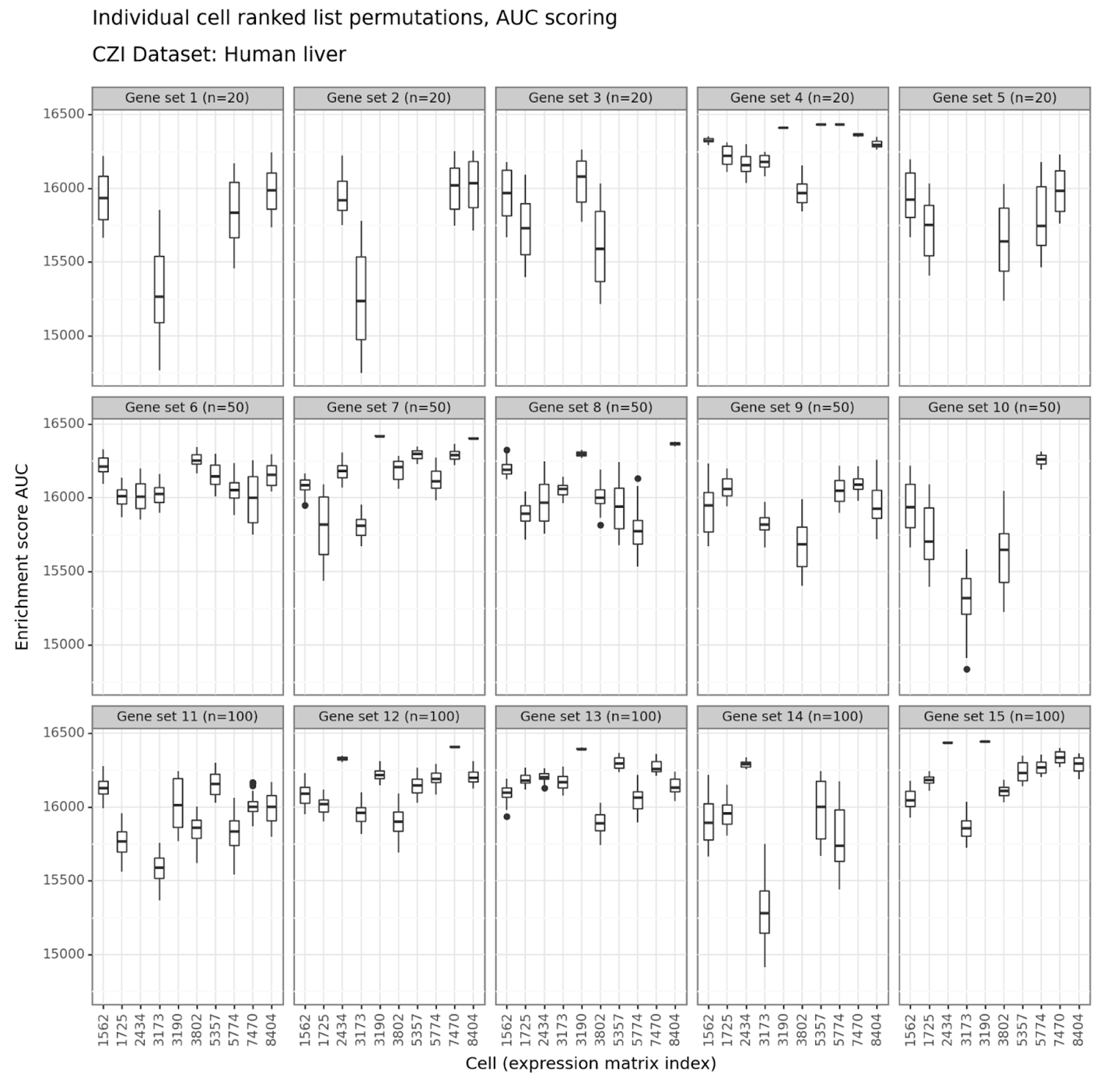

**Figure S7**: Individual cell permutations – Human liver – AUC scores

Figure features as in **Figure 1F**

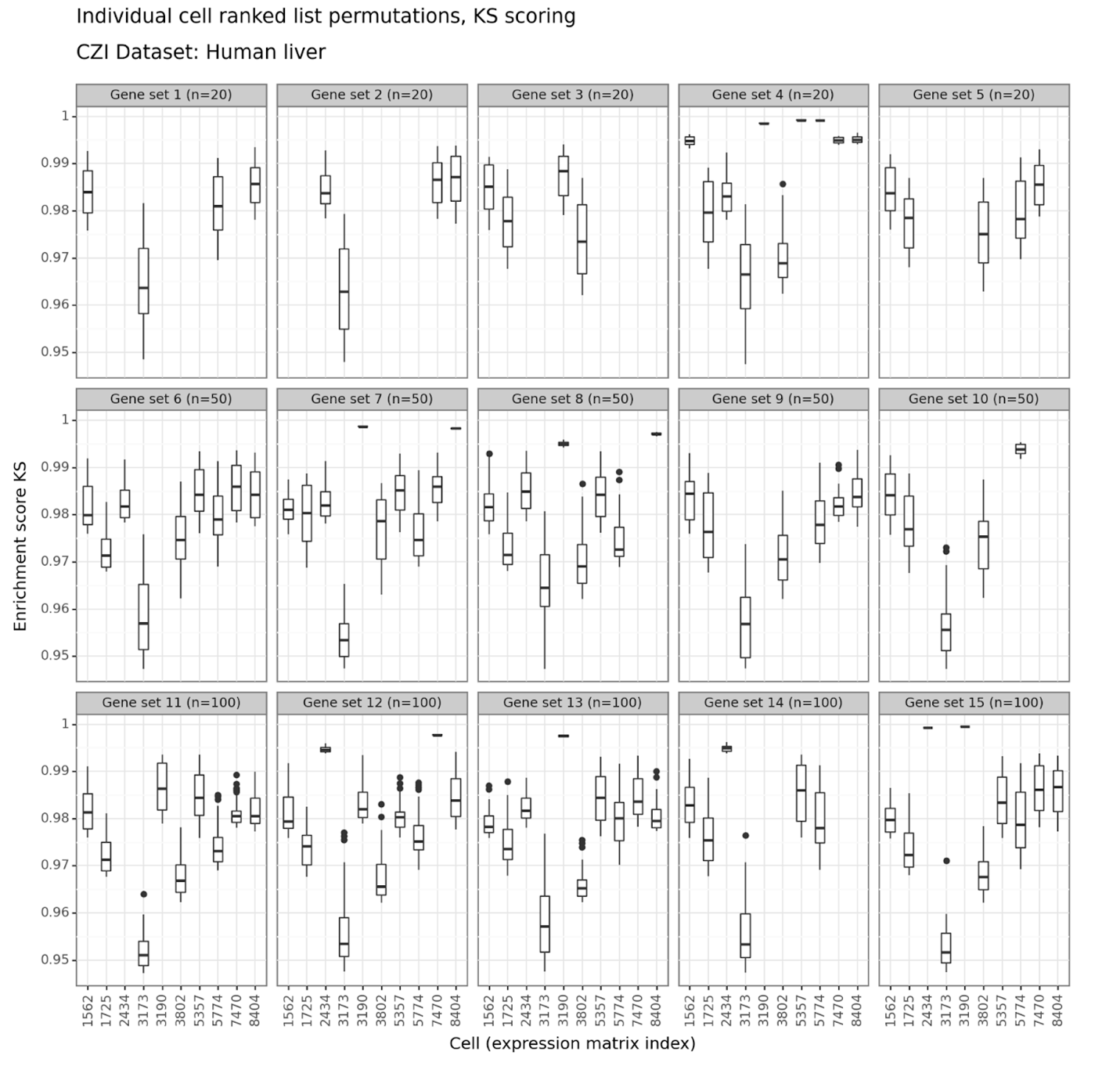

**Figure S8**: Individual cell permutations – Human liver – KS scores

Figure features as in **Figure 1G**

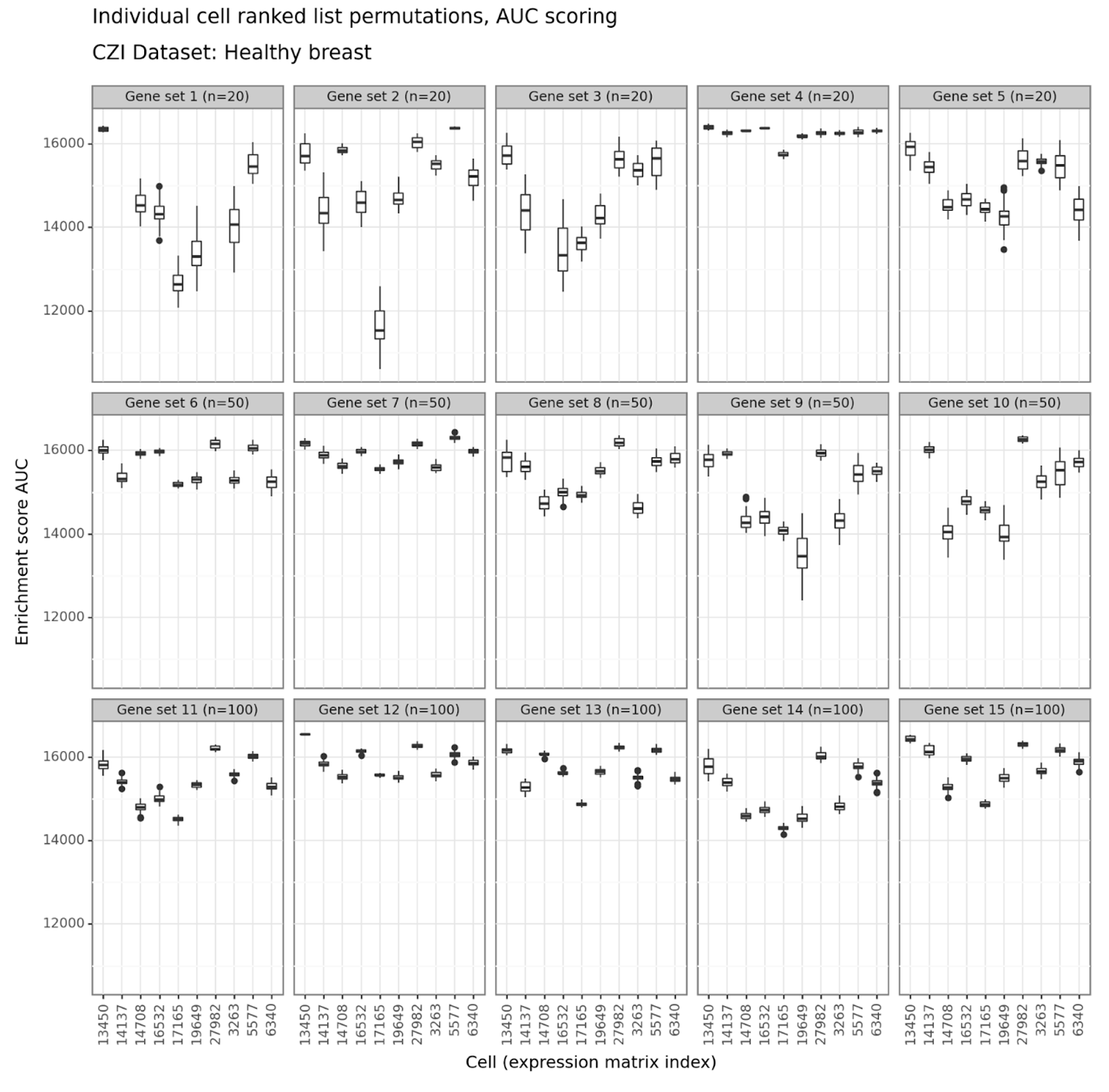

**Figure S9**: Individual cell permutations – Healthy breast – AUC scores

Figure features as in **Figure 1F**

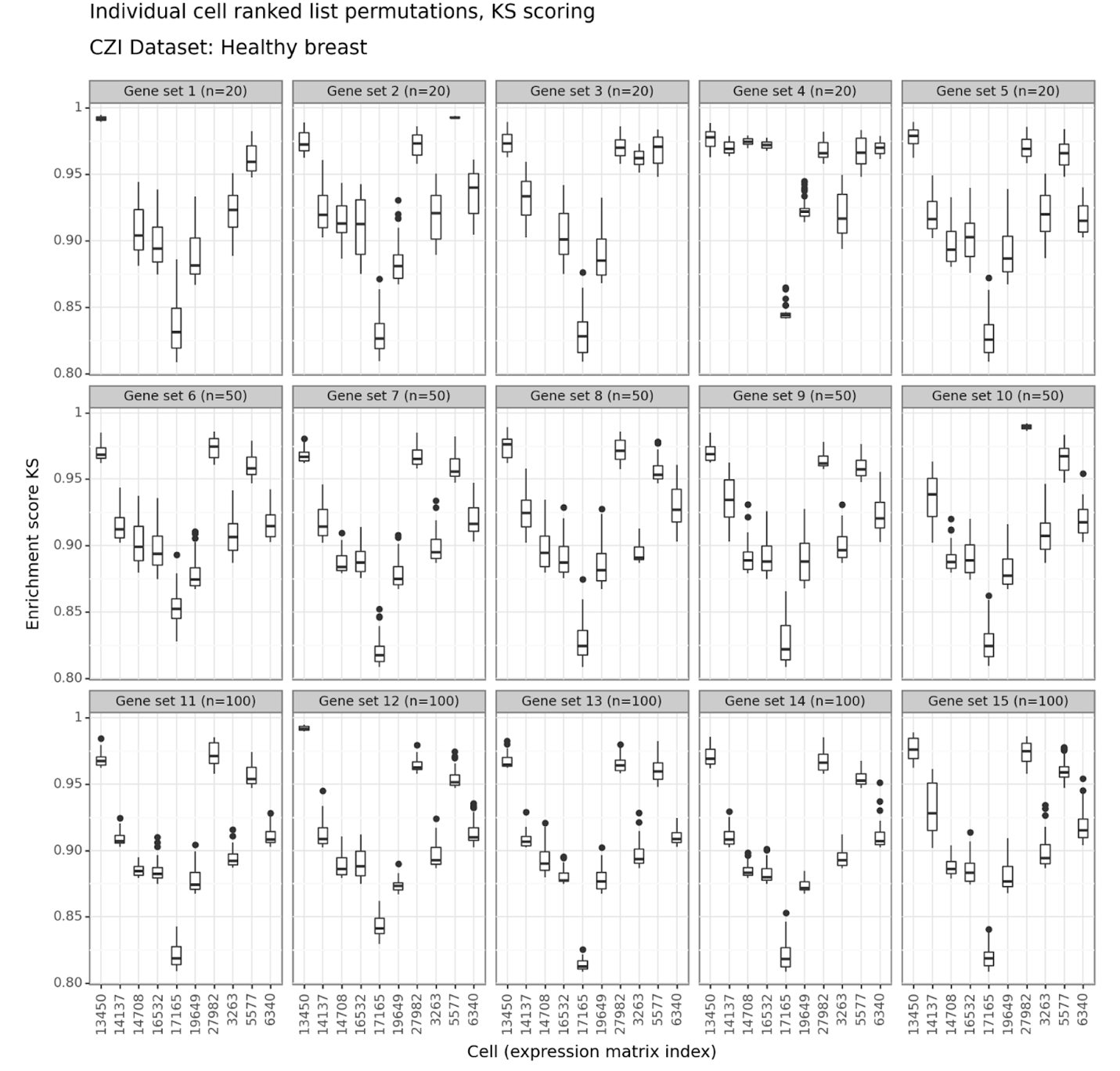

**Figure S10**: Individual cell permutations – Healthy breast – KS scores

Figure features as in **Figure 1G**

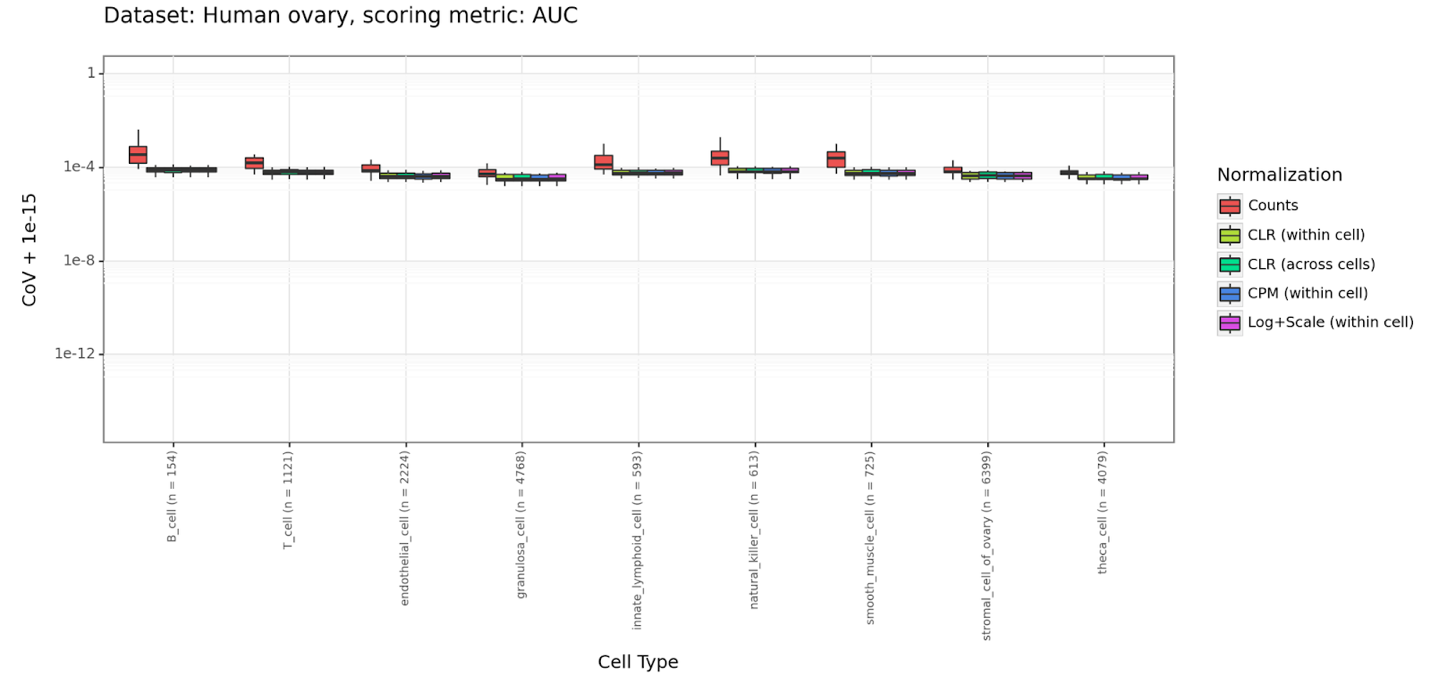

**Figure S11**: Aggregate cell permutations – Human ovary – AUC scores

Figure features as in **Figure 2A**

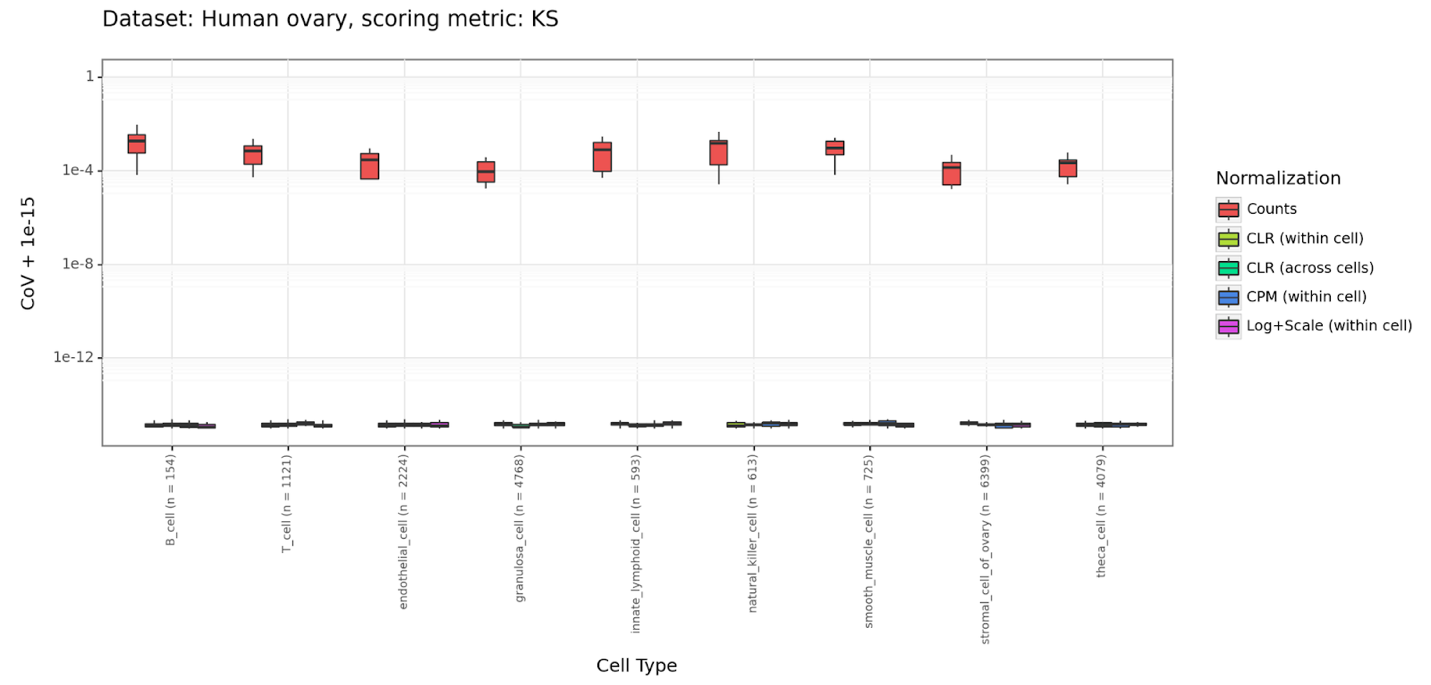

**Figure S12**: Aggregate cell permutations – Human ovary – KS scores

Figure features as in **Figure 2B**

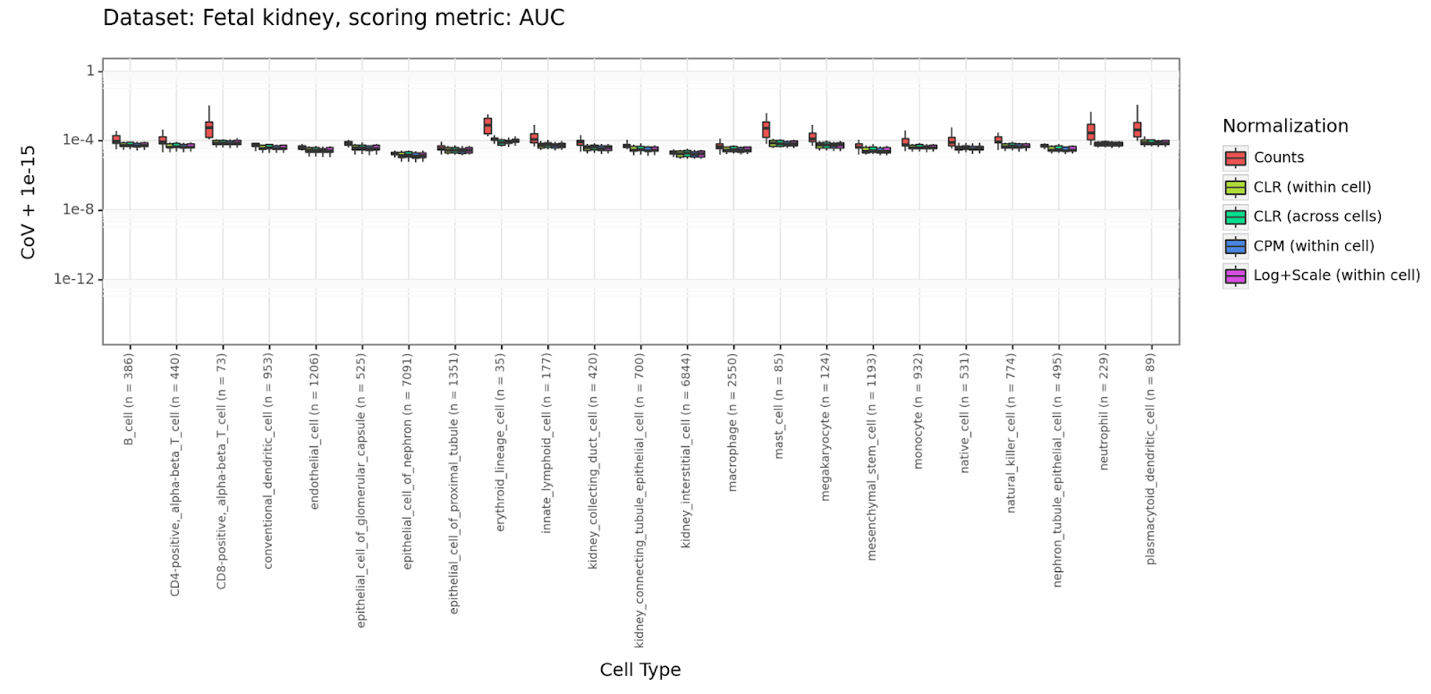

**Figure S13**: Aggregate cell permutations – Fetal kidney – AUC scores

Figure features as in **Figure 2A**

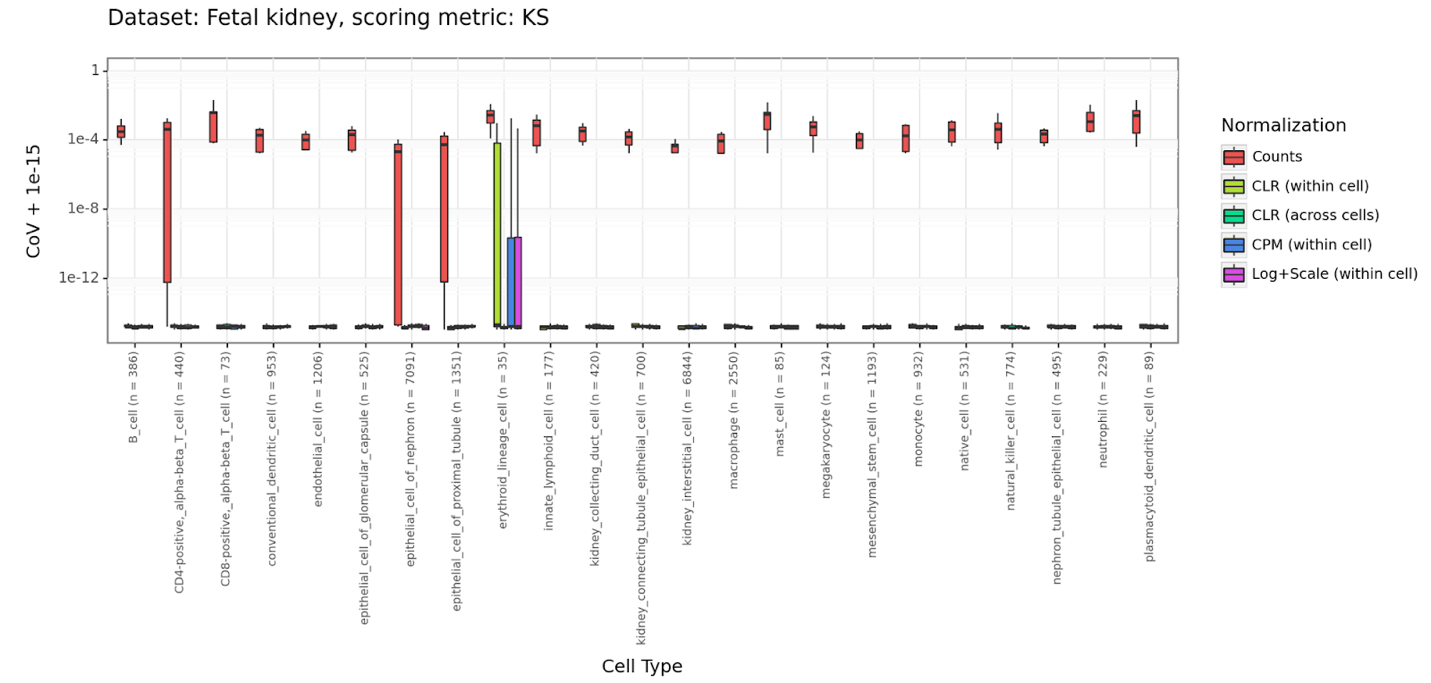

**Figure S14**: Aggregate cell permutations – Fetal kidney – KS scores

Figure features as in **Figure 2B**

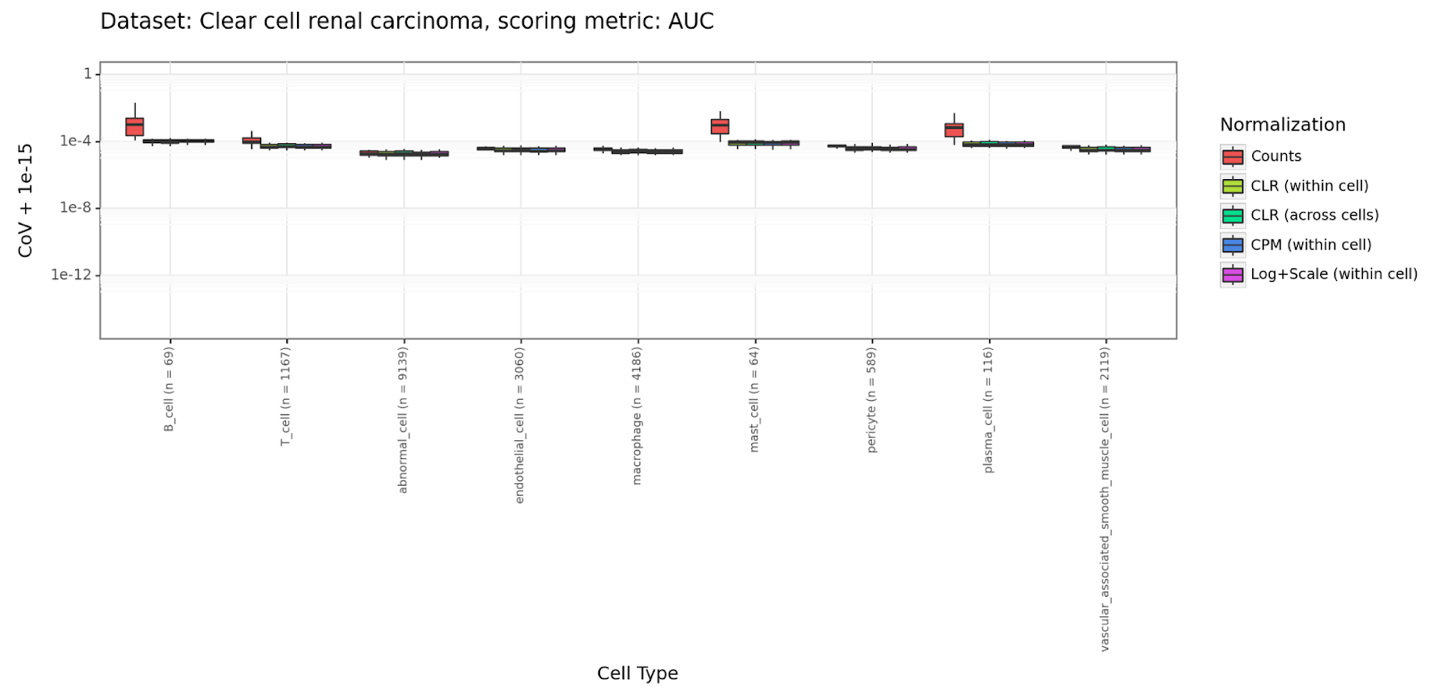

**Figure S15**: Aggregate cell permutations – Clear cell renal carcinoma – AUC scores

Figure features as in **Figure 2A**

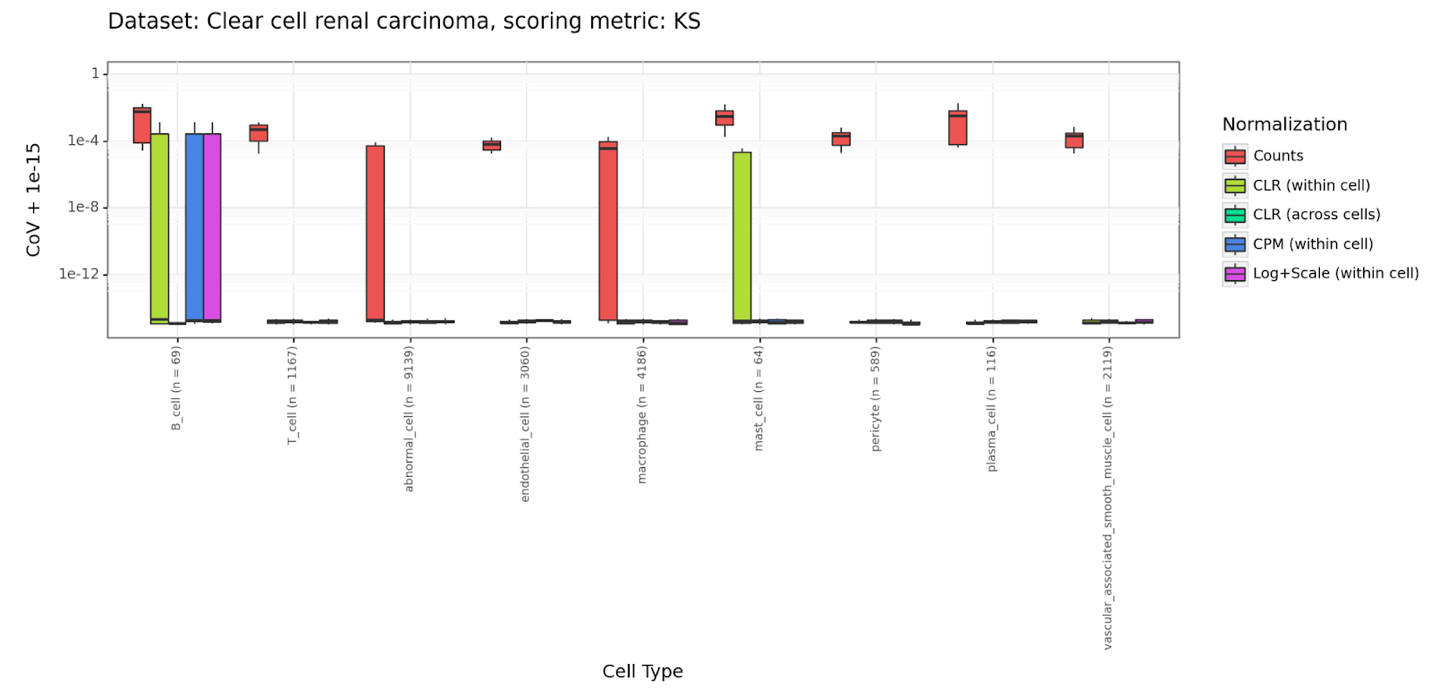

**Figure S16**: Aggregate cell permutations – Clear cell renal carcinoma – KS scores

Figure features as in **Figure 2B**

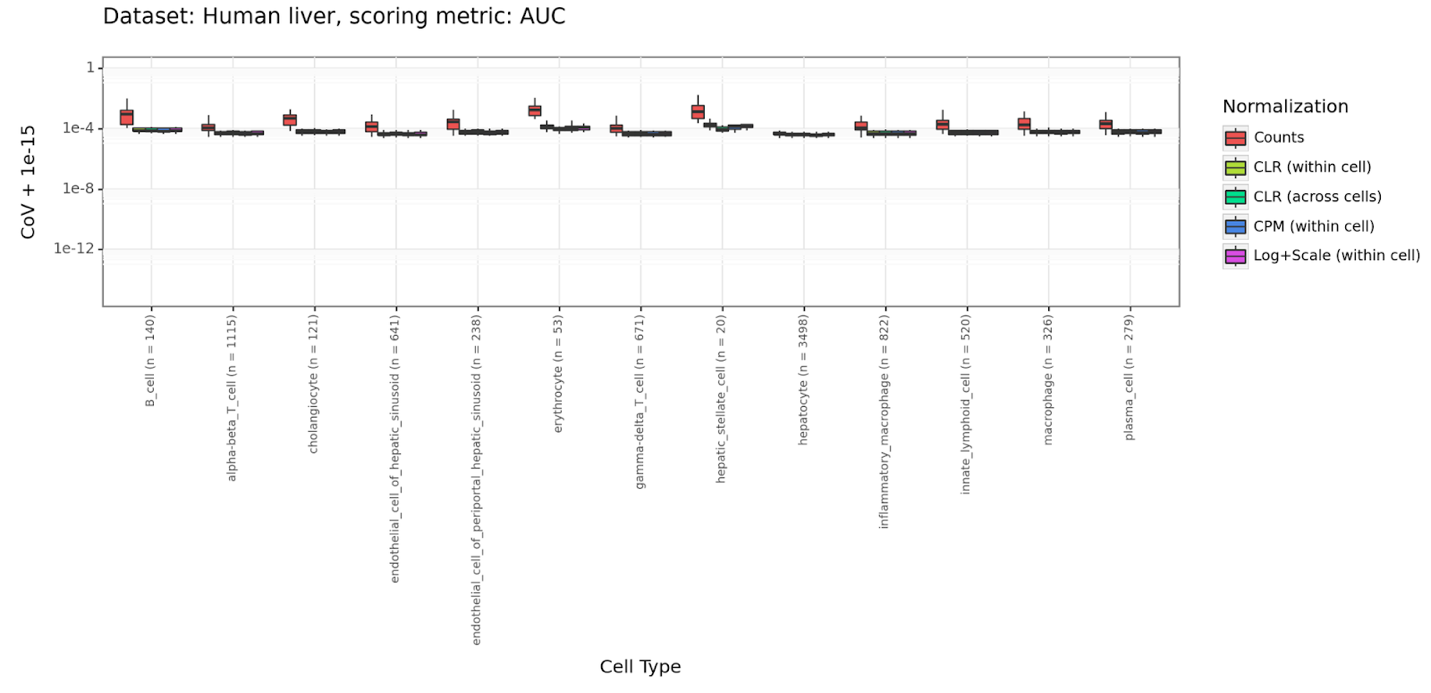

**Figure S17**: Aggregate cell permutations – Human liver – AUC scores

Figure features as in **Figure 2A**

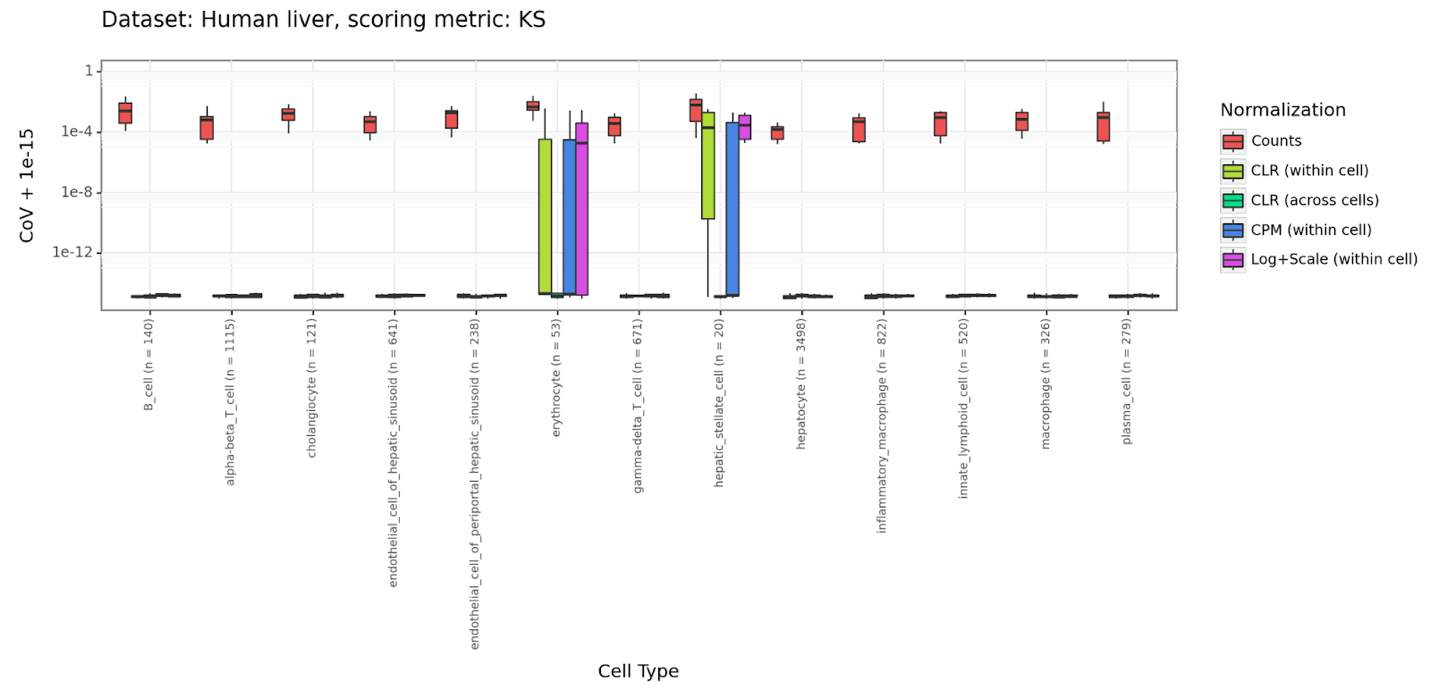

**Figure S18**: Aggregate cell permutations – Human liver – KS scores

Figure features as in **Figure 2B**

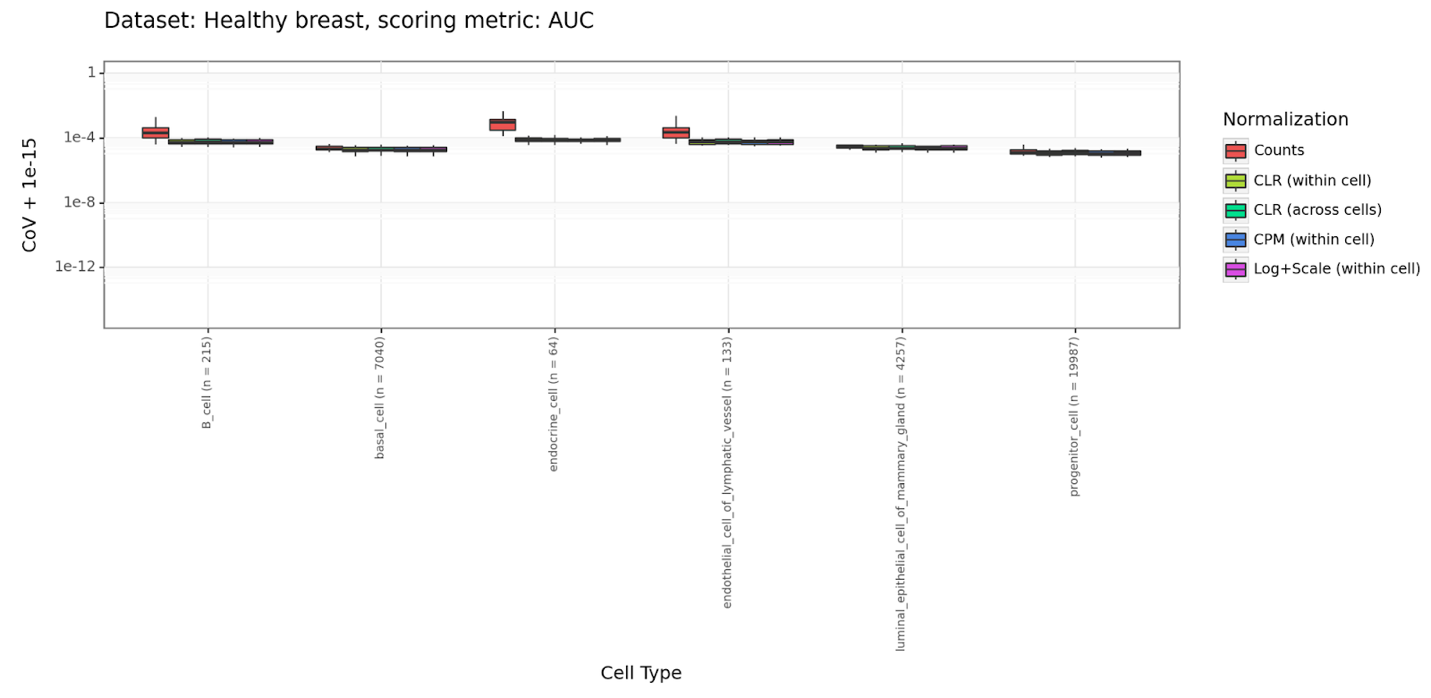

**Figure S19**: Aggregate cell permutations – Healthy breast – AUC scores

Figure features as in **Figure 2A**

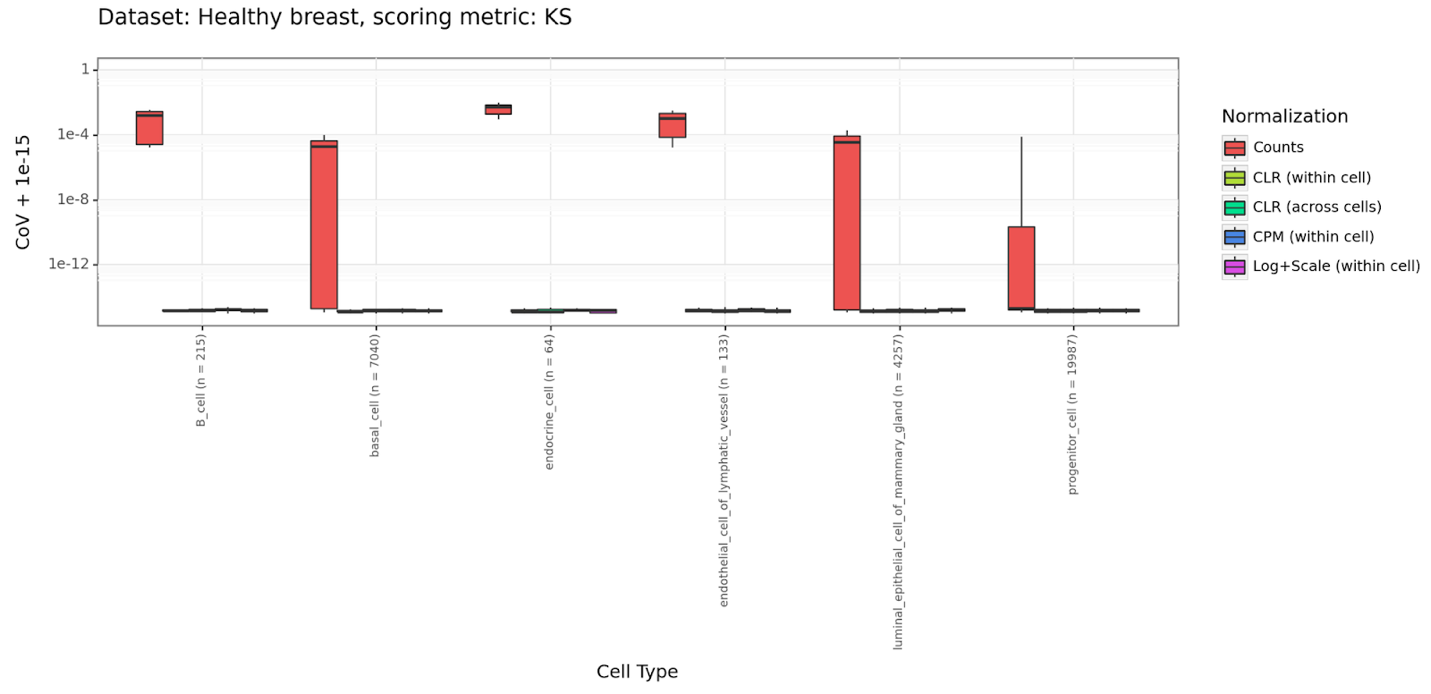

**Figure S20**: Aggregate cell permutations – Healthy breast – KS scores

Figure features as in **Figure 2B**

**Table S2**: Datasets used in individual and aggregate cell benchmarking.

First three columns contain metadata from the CZI CELLxGENE data portal as provided by the authors of the corresponding studies. The fourth column “Brief names” were assigned for the purpose of this work.

| CELLxGENE Dataset ID | CELLxGENE URL | CELLxGENE Full title | Brief name |
| --- | --- | --- | --- |
| 793e90d3-62ab-44c4-98f5-6a090f321ba4 | https://cellxgene.cziscience.com/collections/c353707f-09a4-4f12-92a0-cb741e57e5f0 | Single-cell transcriptomes of the human skin reveal age-related loss of fibroblast priming | Aging Skin |
| e62ae178-63f8-4025-91b2-33b15b48abe3 | https://cellxgene.cziscience.com/collections/2902f08c-f83c-470e-a541-e463e25e5058 | Single-cell reconstruction of follicular remodeling in the human adult ovary | Human ovary |
| 837599ec-6580-42b4-86b7-2fb9ad232734 | https://cellxgene.cziscience.com/collections/13d1c580-4b17-4b2e-85c4-75b36917413f | Single cell derived mRNA signals across human kidney tumors | Fetal kidney |
| 231d025d-6b31-40da-aa38-cf618d53b544 | https://cellxgene.cziscience.com/collections/1df8c90d-d299-4b2e-a54d-a5a80f36e780 | ccRCC - Single-cell analyses of renal cell cancers reveal insights into tumor microenvironment, cell of origin, and therapy response | Clear cell renal carcinoma |
| 91a16284-28c5-41f8-a8c8-b283d2e5b6ea | https://cellxgene.cziscience.com/collections/bd5230f4-cd76-4d35-9ee5-89b3e7475659 | Single cell RNA sequencing of human liver reveals distinct intrahepatic macrophage populations | Human liver |
| 64f14a2b-d754-4bc9-b496-b26f05ebfe4e | https://cellxgene.cziscience.com/collections/c9706a92-0e5f-46c1-96d8-20e42467f287 | A single-cell atlas of the healthy breast tissues reveals clinically relevant clusters of breast epithelial cells | Healthy breast |

**Table S3**: Randomly created gene sets used in benchmarking

| **Gene set name** | **Gene set genes** |
| --- | --- |
| gset_0 | ENSG00000180316, ENSG00000158955, ENSG00000182583, ENSG00000139131, ENSG00000158716, ENSG00000223800, ENSG00000259124, ENSG00000205929, ENSG00000229924, ENSG00000227260, ENSG00000265100, ENSG00000225206, ENSG00000256671, ENSG00000143570, ENSG00000144843, ENSG00000155087, ENSG00000236790, ENSG00000156453, ENSG00000260442, ENSG00000255026 |
| gset_1 | ENSG00000171960, ENSG00000101191, ENSG00000165816, ENSG00000258107, ENSG00000249000, ENSG00000139620, ENSG00000107862, ENSG00000162753, ENSG00000184897, ENSG00000171492, ENSG00000176040, ENSG00000260162, ENSG00000261717, ENSG00000155827, ENSG00000183134, ENSG00000164458, ENSG00000237441, ENSG00000259129, ENSG00000233256, ENSG00000147592 |
| gset_2 | ENSG00000095932, ENSG00000226386, ENSG00000214360, ENSG00000073578, ENSG00000231987, ENSG00000155903, ENSG00000157077, ENSG00000135972, ENSG00000265907, ENSG00000167371, ENSG00000066248, ENSG00000185899, ENSG00000105671, ENSG00000249236, ENSG00000237419, ENSG00000268129, ENSG00000186297, ENSG00000088320, ENSG00000272456, ENSG00000250626 |
| gset_3 | ENSG00000185347, ENSG00000261129, ENSG00000204084, ENSG00000254344, ENSG00000108100, ENSG00000241370, ENSG00000268854, ENSG00000133112, ENSG00000104131, ENSG00000205186, ENSG00000237001, ENSG00000221909, ENSG00000227518, ENSG00000117691, ENSG00000083828, ENSG00000182095, ENSG00000267004, ENSG00000160867, ENSG00000186860, ENSG00000248393 |
| gset_4 | ENSG00000004848, ENSG00000063761, ENSG00000257746, ENSG00000188037, ENSG00000081320, ENSG00000273413, ENSG00000108961, ENSG00000141002, ENSG00000272861, ENSG00000100418, ENSG00000250855, ENSG00000234380, ENSG00000136939, ENSG00000249494, ENSG00000135899, ENSG00000170579, ENSG00000188803, ENSG00000112667, ENSG00000171862, ENSG00000143479 |

| **Gene set name** | **Gene set genes** |
| --- | --- |
| gset_5 | ENSG00000113739, ENSG00000137824, ENSG00000261360, ENSG00000266947, ENSG00000213585, ENSG00000119778, ENSG00000232298, ENSG00000068079, ENSG00000231914, ENSG00000260456, ENSG00000232555, ENSG00000266696, ENSG00000121274, ENSG00000011485, ENSG00000214614, ENSG00000171970, ENSG00000164815, ENSG00000231046, ENSG00000130640, ENSG00000267360, ENSG00000169413, ENSG00000136270, ENSG00000187033, ENSG00000101251, ENSG00000164323, ENSG00000273442, ENSG00000186001, ENSG00000115053, ENSG00000248262, ENSG00000067445, ENSG00000248243, ENSG00000138207, ENSG00000100138, ENSG00000162494, ENSG00000106128, ENSG00000260642, ENSG00000167447, ENSG00000105085, ENSG00000269307, ENSG00000104218, ENSG00000256084, ENSG00000197991, ENSG00000273211, ENSG00000256137, ENSG00000264575, ENSG00000267219, ENSG00000159182, ENSG00000129932, ENSG00000223569, ENSG00000136100 |
| gset_6 | ENSG00000188385, ENSG00000196562, ENSG00000040275, ENSG00000103067, ENSG00000163431, ENSG00000124508, ENSG00000256894, ENSG00000111832, ENSG00000198947, ENSG00000198673, ENSG00000105254, ENSG00000168509, ENSG00000248809, ENSG00000247775, ENSG00000225867, ENSG00000185347, ENSG00000163220, ENSG00000172840, ENSG00000166780, ENSG00000260416, ENSG00000182776, ENSG00000260430, ENSG00000226438, ENSG00000074660, ENSG00000224347, ENSG00000180957, ENSG00000001497, ENSG00000137726, ENSG00000236869, ENSG00000255462, ENSG00000108107, ENSG00000163637, ENSG00000221938, ENSG00000085721, ENSG00000197498, ENSG00000026103, ENSG00000165660, ENSG00000225697, ENSG00000197641, ENSG00000086015, ENSG00000233848, ENSG00000130695, ENSG00000139914, ENSG00000255845, ENSG00000148824, ENSG00000266696, ENSG00000160216, ENSG00000149679, ENSG00000160131, ENSG00000221869 |
| gset_7 | ENSG00000135454, ENSG00000234244, ENSG00000227252, ENSG00000249388, ENSG00000253715, ENSG00000197172, ENSG00000187010, ENSG00000137824, ENSG00000250313, ENSG00000164920, ENSG00000257365, ENSG00000187566, ENSG00000233397, ENSG00000106069, ENSG00000167711, ENSG00000236914, ENSG00000268205, ENSG00000127337, ENSG00000134852, ENSG00000141696, ENSG00000244953, ENSG00000154359, ENSG00000240498, ENSG00000108559, ENSG00000183035, ENSG00000100307, ENSG00000135547, ENSG00000270249, ENSG00000134716, ENSG00000131669, ENSG00000196459, ENSG00000259635, ENSG00000260701, ENSG00000253955, ENSG00000257341, ENSG00000249693, ENSG00000256417, ENSG00000138778, ENSG00000148942, ENSG00000272384, ENSG00000260185, ENSG00000231851, ENSG00000143252, ENSG00000154723, ENSG00000170961, ENSG00000261090, ENSG00000044090, ENSG00000146910, ENSG00000232645, ENSG00000115084 |

| **Gene set name** | **Gene set genes** |
| --- | --- |
| gset_8 | ENSG00000080603, ENSG00000113924, ENSG00000172375, ENSG00000112273, ENSG00000100605, ENSG00000197993, ENSG00000272502, ENSG00000233214, ENSG00000267224, ENSG00000248559, ENSG00000141639, ENSG00000267284, ENSG00000269243, ENSG00000235168, ENSG00000111536, ENSG00000235431, ENSG00000184432, ENSG00000265643, ENSG00000178821, ENSG00000257155, ENSG00000262791, ENSG00000133030, ENSG00000169062, ENSG00000116478, ENSG00000267152, ENSG00000183621, ENSG00000128536, ENSG00000158480, ENSG00000249392, ENSG00000258749, ENSG00000260278, ENSG00000223572, ENSG00000142166, ENSG00000165724, ENSG00000261334, ENSG00000139793, ENSG00000254968, ENSG00000150977, ENSG00000261749, ENSG00000198003, ENSG00000111863, ENSG00000146830, ENSG00000197061, ENSG00000234390, ENSG00000272139, ENSG00000180543, ENSG00000125107, ENSG00000129596, ENSG00000167633, ENSG00000143363 |
| gset_9 | ENSG00000224916, ENSG00000227507, ENSG00000253682, ENSG00000046653, ENSG00000177000, ENSG00000205155, ENSG00000169340, ENSG00000205208, ENSG00000268120, ENSG00000041880, ENSG00000140564, ENSG00000113810, ENSG00000187627, ENSG00000249096, ENSG00000254580, ENSG00000111684, ENSG00000258345, ENSG00000170473, ENSG00000139656, ENSG00000119913, ENSG00000250098, ENSG00000206105, ENSG00000135378, ENSG00000249637, ENSG00000106524, ENSG00000255733, ENSG00000197283, ENSG00000162825, ENSG00000232252, ENSG00000226476, ENSG00000116774, ENSG00000254006, ENSG00000152192, ENSG00000240012, ENSG00000236032, ENSG00000130377, ENSG00000185933, ENSG00000134627, ENSG00000186326, ENSG00000233365, ENSG00000127920, ENSG00000231212, ENSG00000182824, ENSG00000249988, ENSG00000234306, ENSG00000125820, ENSG00000260570, ENSG00000164236, ENSG00000188725, ENSG00000163933 |

| **Gene set name** | **Gene set genes** |
| --- | --- |
| gset_10 | ENSG00000077044, ENSG00000183484, ENSG00000261286, ENSG00000163564, ENSG00000250131, ENSG00000214285, ENSG00000131966, ENSG00000219438, ENSG00000261433, ENSG00000235601, ENSG00000158850, ENSG00000253932, ENSG00000110436, ENSG00000253792, ENSG00000125450, ENSG00000227269, ENSG00000101347, ENSG00000006210, ENSG00000180008, ENSG00000269343, ENSG00000225187, ENSG00000250765, ENSG00000224405, ENSG00000176402, ENSG00000250298, ENSG00000225785, ENSG00000106701, ENSG00000253708, ENSG00000236740, ENSG00000151726, ENSG00000261367, ENSG00000171951, ENSG00000254369, ENSG00000176984, ENSG00000181222, ENSG00000236914, ENSG00000119707, ENSG00000170876, ENSG00000125149, ENSG00000150051, ENSG00000179630, ENSG00000148926, ENSG00000255302, ENSG00000239453, ENSG00000174718, ENSG00000171435, ENSG00000075643, ENSG00000223631, ENSG00000169813, ENSG00000254587, ENSG00000260930, ENSG00000168303, ENSG00000260854, ENSG00000273419, ENSG00000182993, ENSG00000225807, ENSG00000099721, ENSG00000144283, ENSG00000267909, ENSG00000230159, ENSG00000165732, ENSG00000249236, ENSG00000168772, ENSG00000253344, ENSG00000265799, ENSG00000243896, ENSG00000203485, ENSG00000223725, ENSG00000177463, ENSG00000121390, ENSG00000248813, ENSG00000008196, ENSG00000233538, ENSG00000243903, ENSG00000198542, ENSG00000231238, ENSG00000170248, ENSG00000151366, ENSG00000253653, ENSG00000225612, ENSG00000246662, ENSG00000100412, ENSG00000259376, ENSG00000055332, ENSG00000171617, ENSG00000131115, ENSG00000258804, ENSG00000135828, ENSG00000174132, ENSG00000173705, ENSG00000231873, ENSG00000267325, ENSG00000182156, ENSG00000223601, ENSG00000162881, ENSG00000250328, ENSG00000261648, ENSG00000172534, ENSG00000086991, ENSG00000055955 |

| **Gene set name** | **Gene set genes** |
| --- | --- |
| gset_11 | ENSG00000132329, ENSG00000185009, ENSG00000221989, ENSG00000269053, ENSG00000230433, ENSG00000224101, ENSG00000128346, ENSG00000005022, ENSG00000183066, ENSG00000234068, ENSG00000253673, ENSG00000227297, ENSG00000111727, ENSG00000273210, ENSG00000250614, ENSG00000005884, ENSG00000099377, ENSG00000063169, ENSG00000226149, ENSG00000258935, ENSG00000261766, ENSG00000172340, ENSG00000233760, ENSG00000184220, ENSG00000261307, ENSG00000165323, ENSG00000237101, ENSG00000151923, ENSG00000101417, ENSG00000159263, ENSG00000162458, ENSG00000235497, ENSG00000189280, ENSG00000254039, ENSG00000229483, ENSG00000187944, ENSG00000116922, ENSG00000198521, ENSG00000204421, ENSG00000135945, ENSG00000126882, ENSG00000254163, ENSG00000250067, ENSG00000253309, ENSG00000228799, ENSG00000229307, ENSG00000156642, ENSG00000169174, ENSG00000154263, ENSG00000269416, ENSG00000125611, ENSG00000166917, ENSG00000224361, ENSG00000169548, ENSG00000039600, ENSG00000205238, ENSG00000231662, ENSG00000182776, ENSG00000127989, ENSG00000103066, ENSG00000189056, ENSG00000263684, ENSG00000234184, ENSG00000206559, ENSG00000139797, ENSG00000145817, ENSG00000165983, ENSG00000174738, ENSG00000101336, ENSG00000231424, ENSG00000081923, ENSG00000115109, ENSG00000172724, ENSG00000259351, ENSG00000154027, ENSG00000205669, ENSG00000127995, ENSG00000088992, ENSG00000253821, ENSG00000148300, ENSG00000149452, ENSG00000230172, ENSG00000268804, ENSG00000235819, ENSG00000257986, ENSG00000162244, ENSG00000205642, ENSG00000171863, ENSG00000215784, ENSG00000244558, ENSG00000221855, ENSG00000127561, ENSG00000157653, ENSG00000128654, ENSG00000175198, ENSG00000088832, ENSG00000258325, ENSG00000102384, ENSG00000166090, ENSG00000117600 |

| **Gene set name** | **Gene set genes** |
| --- | --- |
| gset_12 | ENSG00000256072, ENSG00000111886, ENSG00000173825, ENSG00000227475, ENSG00000183908, ENSG00000058673, ENSG00000178803, ENSG00000197381, ENSG00000255944, ENSG00000247993, ENSG00000228072, ENSG00000166343, ENSG00000248397, ENSG00000126947, ENSG00000152760, ENSG00000121853, ENSG00000264595, ENSG00000124172, ENSG00000173890, ENSG00000185532, ENSG00000062096, ENSG00000169862, ENSG00000015676, ENSG00000242221, ENSG00000183035, ENSG00000105404, ENSG00000257474, ENSG00000087266, ENSG00000236437, ENSG00000239900, ENSG00000114054, ENSG00000125170, ENSG00000108950, ENSG00000236104, ENSG00000120327, ENSG00000087111, ENSG00000258444, ENSG00000251417, ENSG00000012660, ENSG00000166106, ENSG00000130255, ENSG00000088002, ENSG00000116005, ENSG00000160801, ENSG00000249846, ENSG00000162755, ENSG00000172059, ENSG00000114745, ENSG00000260658, ENSG00000184897, ENSG00000254528, ENSG00000155719, ENSG00000135773, ENSG00000132603, ENSG00000107672, ENSG00000130182, ENSG00000255686, ENSG00000259383, ENSG00000146666, ENSG00000224857, ENSG00000240006, ENSG00000185761, ENSG00000226741, ENSG00000250056, ENSG00000229275, ENSG00000205279, ENSG00000164023, ENSG00000231143, ENSG00000176840, ENSG00000069020, ENSG00000173451, ENSG00000164853, ENSG00000138472, ENSG00000005448, ENSG00000224832, ENSG00000138623, ENSG00000116984, ENSG00000214575, ENSG00000228351, ENSG00000246695, ENSG00000075914, ENSG00000197993, ENSG00000198169, ENSG00000268173, ENSG00000224505, ENSG00000145721, ENSG00000251218, ENSG00000271011, ENSG00000149571, ENSG00000229928, ENSG00000102174, ENSG00000214940, ENSG00000266933, ENSG00000162761, ENSG00000223430, ENSG00000138074, ENSG00000170310, ENSG00000253479, ENSG00000140043, ENSG00000231420 |

| **Gene set name** | **Gene set genes** |
| --- | --- |
| gset_13 | ENSG00000145794, ENSG00000171189, ENSG00000163714, ENSG00000232057, ENSG00000189180, ENSG00000267644, ENSG00000147121, ENSG00000136802, ENSG00000132510, ENSG00000140157, ENSG00000174485, ENSG00000232613, ENSG00000163295, ENSG00000183246, ENSG00000266202, ENSG00000131653, ENSG00000272760, ENSG00000250375, ENSG00000130054, ENSG00000234943, ENSG00000272338, ENSG00000155962, ENSG00000250822, ENSG00000124092, ENSG00000196498, ENSG00000175334, ENSG00000223960, ENSG00000078900, ENSG00000119333, ENSG00000250091, ENSG00000164080, ENSG00000114520, ENSG00000228421, ENSG00000214872, ENSG00000272647, ENSG00000198129, ENSG00000198685, ENSG00000129028, ENSG00000144366, ENSG00000171401, ENSG00000224596, ENSG00000186924, ENSG00000236543, ENSG00000237741, ENSG00000153266, ENSG00000113070, ENSG00000168306, ENSG00000261431, ENSG00000114656, ENSG00000148677, ENSG00000214432, ENSG00000088038, ENSG00000231226, ENSG00000260141, ENSG00000251573, ENSG00000253474, ENSG00000186509, ENSG00000171130, ENSG00000237781, ENSG00000170871, ENSG00000250081, ENSG00000125686, ENSG00000204677, ENSG00000234396, ENSG00000263159, ENSG00000164284, ENSG00000155307, ENSG00000149196, ENSG00000100387, ENSG00000249662, ENSG00000224903, ENSG00000140057, ENSG00000149532, ENSG00000227482, ENSG00000163626, ENSG00000261121, ENSG00000173068, ENSG00000138615, ENSG00000052344, ENSG00000255054, ENSG00000250719, ENSG00000141646, ENSG00000166869, ENSG00000156968, ENSG00000167081, ENSG00000229278, ENSG00000166454, ENSG00000123810, ENSG00000273011, ENSG00000213015, ENSG00000260362, ENSG00000231597, ENSG00000091106, ENSG00000256969, ENSG00000217555, ENSG00000213420, ENSG00000214940, ENSG00000247708, ENSG00000059573, ENSG00000135318 |

| **Gene set name** | **Gene set genes** |
| --- | --- |
| gset_14 | ENSG00000124532, ENSG00000248235, ENSG00000125246, ENSG00000174292, ENSG00000183423, ENSG00000081760, ENSG00000175065, ENSG00000146828, ENSG00000226067, ENSG00000272259, ENSG00000257135, ENSG00000225243, ENSG00000262358, ENSG00000204278, ENSG00000143811, ENSG00000148343, ENSG00000186910, ENSG00000268810, ENSG00000227851, ENSG00000268754, ENSG00000187754, ENSG00000102934, ENSG00000101190, ENSG00000267726, ENSG00000110025, ENSG00000262133, ENSG00000233651, ENSG00000226516, ENSG00000259529, ENSG00000052795, ENSG00000231128, ENSG00000260470, ENSG00000260303, ENSG00000138115, ENSG00000229671, ENSG00000153086, ENSG00000221923, ENSG00000267204, ENSG00000188859, ENSG00000185668, ENSG00000168496, ENSG00000100162, ENSG00000248508, ENSG00000273149, ENSG00000233234, ENSG00000261448, ENSG00000163629, ENSG00000167526, ENSG00000198001, ENSG00000007372, ENSG00000254136, ENSG00000227059, ENSG00000272502, ENSG00000188828, ENSG00000107036, ENSG00000259523, ENSG00000236668, ENSG00000230552, ENSG00000214970, ENSG00000155886, ENSG00000112837, ENSG00000257634, ENSG00000269901, ENSG00000255548, ENSG00000234906, ENSG00000164076, ENSG00000158806, ENSG00000204228, ENSG00000261227, ENSG00000122952, ENSG00000185352, ENSG00000260088, ENSG00000272431, ENSG00000269292, ENSG00000232767, ENSG00000237837, ENSG00000070729, ENSG00000249924, ENSG00000260759, ENSG00000155897, ENSG00000105971, ENSG00000162408, ENSG00000242082, ENSG00000214413, ENSG00000171819, ENSG00000258580, ENSG00000223991, ENSG00000205863, ENSG00000164841, ENSG00000230894, ENSG00000137947, ENSG00000255372, ENSG00000198874, ENSG00000248973, ENSG00000134538, ENSG00000263688, ENSG00000258853, ENSG00000250049, ENSG00000178700, ENSG00000257835 |
